# M1-linked Ubiquitination by LUBAC Regulates AMPK Activity and the Response to Energetic Stress

**DOI:** 10.1101/2024.11.08.622598

**Authors:** Camilla Reiter Elbæk, Akhee S. Jahan, Joyceline Cuenco, Anna M. Dahlström, Anna L. Aalto, John Rizk, Simon A. Hawley, Sarah N. J. Franks, Michael Stumpe, Srinivasa Prasad Kolapalli, Ximena Hildebrandt, Alexander Erwin Früh, Klara Nielsen, Dominik Priesmann, Julian Koch, Mathilde Deichmann, Josef Gullmets, Lien Verboom, Geert van Loo, Nieves Peltzer, Lisa B. Frankel, Mads Gyrd-Hansen, Jörn Dengjel, Brent J. Ryan, D. Grahame Hardie, Annika Meinander, Kei Sakamoto, Rune Busk Damgaard

**Author notes:** Correspondence: Rune Busk Damgaard.

## Abstract

Methionine-1 (M1)-linked ubiquitin chains, assembled by the ubiquitin ligase LUBAC and cleaved by the deubiquitinase OTULIN, are critical regulators of inflammation and immune homeostasis. Genetic loss of either LUBAC or OTULIN causes autoinflammatory syndromes, which are associated with defects in glycogen and lipid metabolism. However, how LUBAC and OTULIN regulate metabolic signalling remains unknown. Here, we demonstrate that LUBAC promotes, while OTULIN restricts, activation of the key metabolic regulator AMP-activated protein kinase (AMPK) in cells, mice, and human samples. LUBAC and OTULIN interact with AMPK, control its M1-ubiquitination, and regulate its activation in response to glucose starvation and allosteric activation. During starvation, LUBAC deficiency impairs autophagy induction and hinders the shift from oxidative phosphorylation to glycolysis. Strikingly, LUBAC-deficient *Drosophila* have a strongly reduced survival rate after starvation. Our work identifies LUBAC and OTULIN as physiological regulators of AMPK, providing the first mechanism by which M1-linked ubiquitin chains regulate metabolic signalling.

## Introduction

Ubiquitination is a pervasive and versatile post-translational modification that controls a wide range of cellular functions, including proteostasis, autophagic pathways, the DNA damage response, and inflammatory NF-κB signalling, through degradative and non-degradative modifications^1^. Ubiquitin (Ub) mediates most of its cellular functions through structurally and functionally distinct polyUb signals^1,2^. These polyUb chains can be linked via one of the seven Lys (K) residues in Ub (e.g. K48-linked chains) or via Ub Met1 (M1), forming M1-linked (also known as linear) chains^2^.

M1-linked Ub chains are assembled by the linear ubiquitin chain assembly complex (LUBAC), consisting of HOIP, HOIL-1, and SHARPIN^3^, with HOIP serving as the catalytic subunit^4,5^. LUBAC-mediated M1-linked Ub signalling is counteracted by deubiquitinases (DUBs), primarily OTU DUB with linear linkage specificity (OTULIN) and cylindromatosis (CYLD). OTULIN exclusively cleaves M1-linked Ub chains^6,7^, whereas CYLD cleaves both M1- and K63-linked chains^8,9^. Both OTULIN and CYLD interact with LUBAC through the N-terminal PUB domain in HOIP^10–14^. LUBAC is an important regulator of innate immune pathways, particularly TNF signalling, NF-κB activation, and cell death and is recruited to many immune receptors, including TNF Receptor-1 (TNF-R1), in a Ub-dependent manner^3^. At the receptors, LUBAC conjugates non-degradative M1-linked Ub chains to a range of substrates directly or on pre-existing Ub chains, to form a recruitment and activation scaffold for the IκB kinase (IKK) complex, which induces NF-κB activation and drives the inflammatory response^3,15^.

While M1-linked Ub chains constitute only ∼0,5% of all Ub chains in cells^16^, they are essential for maintaining tissue and immune homeostasis. In mice, genetic loss of LUBAC subunits or OTULIN causes embryonic lethality^7,17,18^ or inflammatory phenotypes^19–22^. Loss-of-function mutations in the LUBAC subunits HOIP, HOIL-1, or SHARPIN, or in OTULIN cause early-onset, TNF-mediated, potentially lethal autoinflammatory syndromes in humans, termed LUBAC deficiency and OTULIN-related autoinflammatory syndrome (ORAS), respectively^16,19,23–26^. Both LUBAC deficiency and ORAS are also associated with metabolic dysregulation. HOIP and HOIL-1 mutations cause amylopectinosis, a form of glycogen storage disease with accumulation of unbranched glycogen (polyglucosan), in the heart and skeletal muscle^23,24,27^, whereas OTULIN deficiency is associated with lipodystrophy, subcutaneous fat inflammation (panniculitis), and liver steatosis^16,19,26,28^. In mice, hepatocyte-specific deletion of OTULIN causes severe dysmetabolism, mammalian target of rapamycin complex 1 (mTORC1) dysregulation, and hepatocellular carcinoma in a TNF-R1-independent manner^28,29^. However, it remains unclear which signalling pathways and molecular mechanisms are regulated by LUBAC, OTULIN, and M1-linked Ub chains in controlling metabolism, or how their metabolic control contributes to the clinical manifestations of LUBAC deficiency and ORAS.

Maintaining energy and metabolic homeostasis is vital for cellular function and survival and is achieved through intricate molecular mechanisms that sense and respond to fluctuations in energy and nutrient availability. Central to this regulatory network is AMP-activated protein kinase (AMPK), a highly conserved Ser/Thr kinase activated in response to cellular energy depletion^30^. Due to its vital roles in energy homeostasis and critical physiological functions such as fatty acid and cholesterol synthesis, mitochondrial biogenesis, autophagy, and insulin sensitivity, AMPK is an attractive therapeutic target in various diseases, including type-2 diabetes, metabolic syndrome and fatty liver disease, cancer, and neurodegenerative disorders^31–34^.

AMPK forms a heterotrimeric complex consisting of a catalytic subunit (α) and two regulatory subunits (β and γ)^35^. Canonical AMPK activation results from increases in intracellular AMP:ATP or ADP:ATP ratios, indicating a drop in cellular energy status^36^. AMP or ADP binds to the AMPKγ subunit, displacing ATP, which leads to a conformational rearrangement in AMPK that exposes the conserved Thr172 (T172) residue on the α1 or α2 subunit for phosphorylation by the upstream kinases Liver Kinase B1 (LKB1)^37–39^ or Ca^2+^/Calmodulin-dependent Protein Kinase Kinase 2 (CaMKK2), thereby activating AMPK^40,41^. AMPK is a key metabolic regulator that orchestrates a range of metabolic switches and adaptive responses aimed at restoring energy balance by phosphorylating numerous targets that inhibit anabolism and promote catabolism^30^. AMPK inhibits fatty acid synthesis and activates fatty acid oxidation by phosphorylation of acetyl-CoA carboxylases 1 (ACC1) and 2 (ACC2) respectively^42–44^, inhibits protein synthesis by phosphorylating RAPTOR and TSC2 to inhibit mTORC1^45,46^ or by directly phosphorylating eEF2K^47^, and regulates autophagy either by direct phosphorylation of unc-51-like autophagy activating kinase 1 (ULK1)^48,49^ or through mTORC1 inhibition^50^. Moreover, AMPK activity can modulate inflammatory signalling and is involved in immunometabolic programming of T cells and macrophages^51^. AMPK signalling is known to primarily be regulated by phosphorylation^52^, and the role of ubiquitination in AMPK regulation is not well understood.

Here, we reveal unexpected and important roles of LUBAC, OTULIN, and M1-linked Ub chains in AMPK regulation and control of cellular metabolism and identify M1-linked non-degradative ubiquitination as a new level of regulation of AMPK activation. We show that LUBAC and OTULIN are physiological regulators of AMPK signalling across multiple cell types, tissues, and species, and that LUBAC activity promotes AMPK activation during energetic stress, while OTULIN restricts it. Notably, AMPK activation induced by glucose starvation does not lead to activation of IKK or TBK1 and is not dependent on TNF signalling, showing that the functions of LUBAC and OTULIN in AMPK signalling are unrelated to and independent of their functions in inflammatory signalling. Mechanistically, LUBAC controls both the amplitude and threshold of AMPK activation in response to glucose starvation and allosteric activation, and LUBAC is important for starvation-induced autophagy and the switch from oxidative phosphorylation to glycolysis. Strikingly, LUBAC-deficient *Drosophila* exhibit a profoundly reduced survival rate after starvation. Collectively, this study reveals the first mechanism by which LUBAC and M1-linked Ub directly regulates metabolic signalling and identifies a new level of regulation of AMPK by non-degradative ubiquitination.

## Results

### LUBAC and OTULIN Form a Complex with AMPK in Cells

We previously reported metabolic dysregulation in mice with hepatocyte-specific deletion of OTULIN^28^, but it remains unknown how M1-linked Ub chains regulate metabolism. To identify M1-linked Ub-regulated pathways associated with metabolism, we expressed Strep-tagged OTULIN in HEK293T cells, enriched for OTULIN and its interactors by Strep-Tactin pulldown, and examined the OTULIN interactome using label-free liquid chromatography-tandem mass spectrometry (LC-MS/MS)-based proteomics (**Table S1**). We analysed the interactomes of full-length wild-type (WT) OTULIN and a truncated form of OTULIN with a deletion of its C-terminal PDZ domain-binding motif (ΔPBM) (**Figures 1A and S1A**). The PBM deletion disrupts OTULIN’s interaction with SNX27 and the endosomal retromer trafficking compartment^53^, allowing us to identify SNX27- and retromer-independent interactions. We identified several known interactors of full-length OTULIN, including HOIP (RNF31) and SNX27 (**Figure 1B and Table S1**). As expected, PDZ domain-containing proteins, including SNX27 and SCRIB, were not enriched in the OTULIN^ΔPBM^ interactome (**Figures 1B and S1B**). Interestingly, we identified multiple putative OTULIN interactors in the AMPK signalling pathway, including the catalytic AMPKα1 subunit (PRKAA1) and the AMPK substrates RAPTOR^46^ and TSC2^45^ (**Figure 1B**). The interactions between OTULIN and the AMPK pathway proteins were generally not affected by deletion of the PBM (**Figure 1B**), indicating that they are not indirect interactors from the endosomal trafficking compartment, and suggesting that OTULIN is a putative regulator of AMPK signalling.

**Figure 1.**
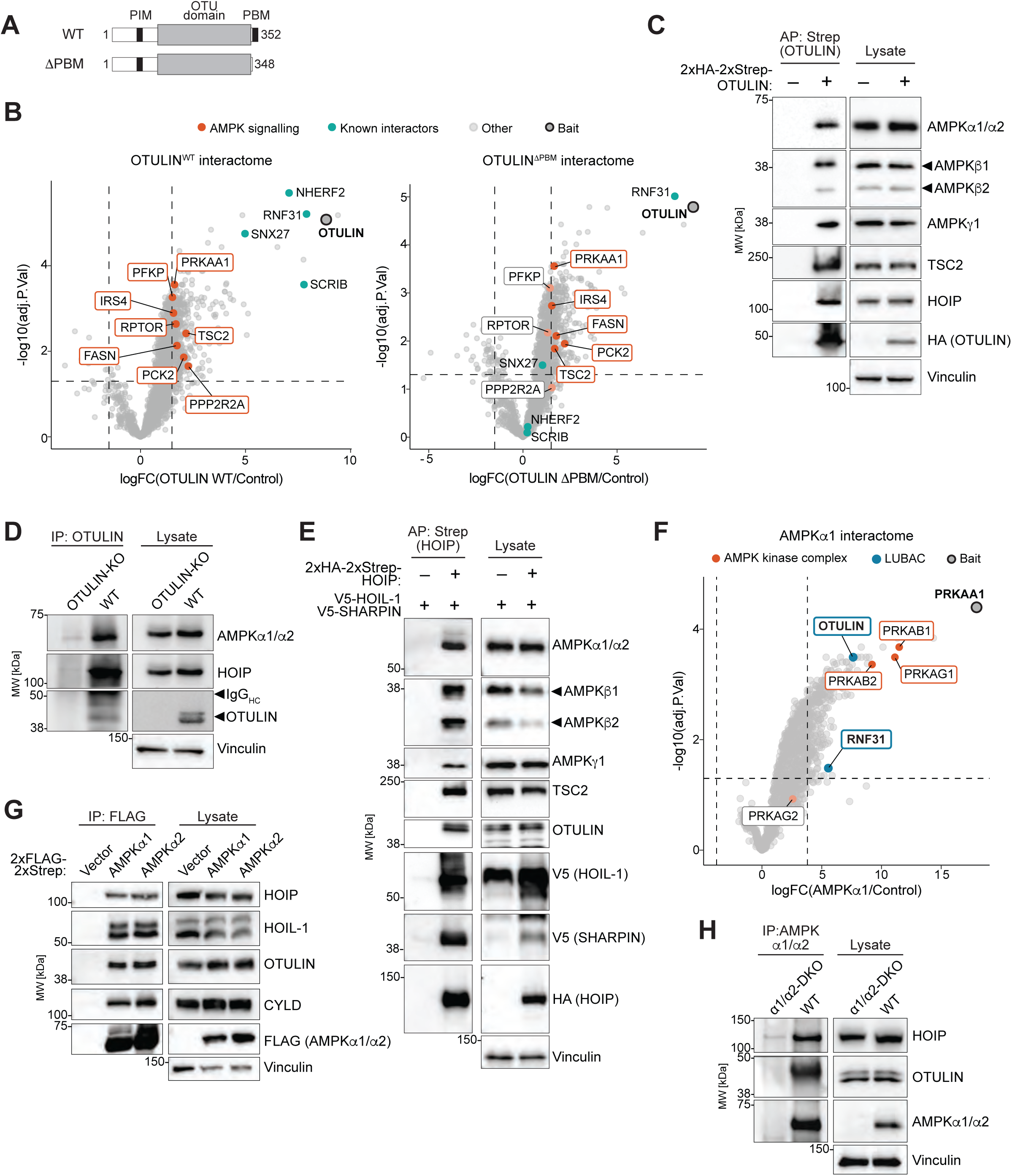
MS-based Proteomics Identify LUBAC and OTULIN as Interactors of AMPK in Cells. A) Schematic domain representation of OTULIN wildtype (WT) and PDZ domain binding motif-deleted (ΔPBM) mutant. B) Volcano plots depicting log2 fold-changes (logFC) of proteins identified in MS analyses of Strep-pulldown from HEK293T cells expressing Strep-tagged OTULIN WT (left) or OTULIN ΔPBM (right) compared with vector control. Known interactors of OTULIN (teal) and significantly enriched proteins within the AMPK signalling pathway (orange) are indicated. BH-adjusted false discovery rate (FDR)<0.05 and a log2 fold-change cutoff of 2.5 were applied. C) Strep-tagged OTULIN was affinity-precipitated (AP) from lysates of HEK293T cells transfected with empty vector or 2xHA-2xStrep-OTULIN. The precipitate was examined for copurifying proteins by immunoblotting. D) Endogenous OTULIN was immunoprecipitated from lysates of HEK293T cells and examined for copurifying proteins by immunoblotting. OTULIN-KO cells were used as IP control. E) Strep-tagged HOIP was affinity-precipitated (AP) from lysates of HEK293T cells transfected with V5-HOIL1, V5-SHARPIN, and 2xHA-2xStrep-HOIP or empty vector control and analysed by immunoblotting. F) Volcano plot showing log2 fold-changes (logFC) of proteins identified in MS analysis of Strep-pulldown from HEK293T cells expressing Strep-tagged AMPKα1 or empty vector. AMPK subunits (orange), as well as OTULIN and HOIP (blue), are indicated. BH-adjusted FDR<0.05 and a log2 fold-change cutoff of 2.5 were applied. G) FLAG-AMPKα1 and FLAG-AMPKα2 were immunoprecipitated from HEK293T cells transfected with empty vector, 2xFLAG-2xStrep-AMPKα1, or 2xFLAG-2xStrep-AMPKα2 and analysed by immunoblotting. H) Endogenous AMPKα1/α2 was immunoprecipitated from lysates of HEK293T cells and examined for copurified proteins by immunoblotting. AMPKα1/α2 double-KO cells were used as IP control. All blots show a single experiment representative of three biologically independent replicates.

We validated the interaction between OTULIN and AMPK in HEK293T cells by expression and affinity purification of Strep-tagged OTULIN. SDS-PAGE followed by immunoblot analysis showed that AMPKα1/2, as well as the β1, β2 and γ1 subunits of AMPK, and the AMPK substrate TSC2 co-precipitated with OTULIN (**Figure 1C**). Endogenous OTULIN was also able to co-precipitate AMPKα1/2 (**Figure 1D**), showing that OTULIN and AMPK interact in cells. Since OTULIN interacts with and antagonises LUBAC, we hypothesised that LUBAC also interacts with AMPK, suggesting a mechanism for AMPK regulation by M1-linked ubiquitination. To test if LUBAC interacted with AMPK, we expressed HOIP, HOIL-1, and SHARPIN in cells and affinity purified LUBAC via Strep-tagged HOIP. Indeed, AMPK α1/2, β1, β2, and γ1 subunits all co-precipitated with LUBAC (**Figure 1E**), confirming the interaction between LUBAC and the heterotrimeric AMPK complex.

To substantiate the novel interactions between LUBAC, OTULIN, and AMPK further, we affinity purified FLAG-tagged AMPKα1 from cells and analysed its interactors by MS-based proteomics (**Figure S1C and Table S2**). The AMPKα1 (PRKAA1) interactome identified both OTULIN and HOIP (RNF31) among the significantly enriched proteins, along with AMPK β1 (PRKAB1), β2 (PRKAB2), and γ1 (PRKAG1) (**Figure 1F**) and more than 100 overlapping interactors between OTULIN^WT^ and AMPK (**Figure S1D**). Immunoprecipitation and immunoblot analysis showed that both FLAG-tagged AMPKα1 and α2 co-precipitated the HOIP and HOIL-1 subunits of LUBAC as well as OTULIN and CYLD (**Figure 1G**), showing that the M1-linked Ub machinery does not discriminate between different AMPK complexes and interacts with complexes containing either catalytic subunit. Finally, endogenous AMPKα1/2 also co-immunoprecipitated both HOIP and OTULIN (**Figure 1H**).

Collectively, these data show that LUBAC, OTULIN, and the key metabolic regulator AMPK form a complex in cells.

### LUBAC and OTULIN Regulate AMPK Signalling in Response to Energetic Stress

The presence of a LUBAC-OTULIN-AMPK complex in cells prompted us to test if LUBAC and OTULIN are important for AMPK signalling. We therefore starved cells for glucose and analysed AMPK signalling by SDS-PAGE and immunoblotting. In U2OS cells, siRNA-mediated knockdown of LUBAC using a mix of siRNAs targeting HOIP, HOIL-1, and SHARPIN, or knockdown of the catalytic subunit HOIP alone, reduced the phosphorylation of AMPKα at T172 in response to glucose starvation for up to six hours compared with control siRNA (**Figures 2A and S2A-B**). Knockdown of LUBAC led to an overall reduction in the amplitude of AMPKα T172 phosphorylation at all timepoints during starvation without overtly changing the kinetics of signalling. The effect of LUBAC knockdown was similar at the level of phosphorylation of the AMPK substrates ACC (S79) and RAPTOR (S792) (**Figures 2A and S2A-B**). We also observed a reduction in starvation-induced AMPK signalling, particularly ACC and RAPTOR phosphorylation, in CRISPR-generated HOIP knockout (KO) cells (**Figure 2B**). Although the effect of permanent HOIP loss in these cells were milder than the effect of transient LUBAC knockdown, impaired AMPK signalling was apparent as reduced ACC and RAPTOR phosphorylation, particularly at 1-4 hours of starvation (**Figure 2B**). This genetically confirmed that HOIP promotes AMPK signalling. Interestingly, we found that the catalytic ubiquitin ligase activity of HOIP, not only the presence of HOIP or LUBAC protein, was critical for promoting AMPK signalling. In mouse embryonic fibroblasts (MEFs) with knock-in of the HOIP^C879S^ mutation, in which HOIP’s catalytic cysteine is replaced by a serine^5^, AMPK, ACC, and RAPTOR phosphorylation was strongly reduced in response to glucose starvation compared with WT MEFs (**Figure 2C**). AMPK is important for metabolic homeostasis in the liver^54–56^. We therefore tested if LUBAC was important for AMPK signalling in a hepatocyte cell line. Consistent with our observations from U2OS cells and MEFs, CRISPR-mediated KO of HOIP in AML12 cells, a non-transformed hepatocyte cell line, markedly reduced the phosphorylation of AMPK, ACC, and RAPTOR in response to glucose starvation (**Figure 2D**). Together, these data show that LUBAC is required for proper and full activation of AMPK pathway signalling in response to glucose starvation in multiple cell types, including cells from metabolic tissues.

**Figure 2.**
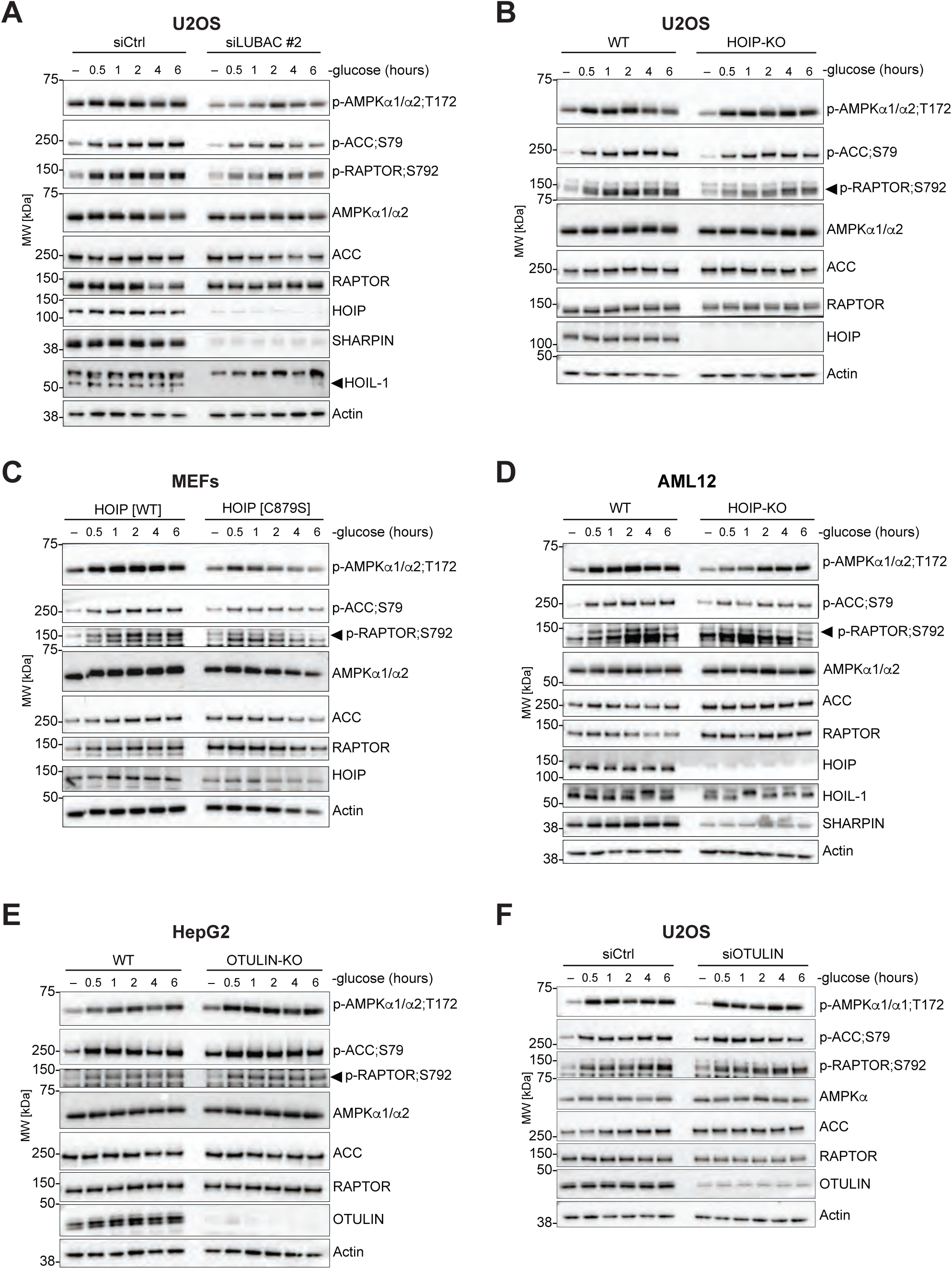
LUBAC and OTULIN Regulate AMPK Signalling After Glucose Starvation. A) U2OS cells were transfected with siRNAs targeting HOIP (oligo #2), HOIL-1, and SHARPIN (siLUBAC #2) or non-targeting control (siCtrl), starved in glucose-free medium as indicated, and analysed by immunoblotting. B) U2OS WT and HOIP-KO cells were treated and analysed as in (A). C) WT or knock-in mouse embryonic fibroblasts (MEFs) expressing catalytically inactive HOIP-C879S were treated and analysed as in (A). D) AML12 WT and HOIP KO cells were treated and analysed as in (A). E) HepG2 WT and OTULIN KO cells were treated and analysed as in (A). F) U2OS cells transfected with siRNAs targeting OTULIN or siCtrl were treated and analysed as in (A). All blots show a single experiment representative of three biologically independent replicates.

OTULIN antagonises LUBAC and restricts M1-linked Ub signalling in the NF-κB pathway and in inflammation^6,7,19,57^. To investigate if OTULIN also restricted AMPK signalling, we starved WT and OTULIN-KO HepG2 hepatocytes of glucose. OTULIN-deficient HepG2 cells displayed enhanced AMPK signalling with increased AMPK, ACC and RAPTOR phosphorylation, in response to glucose withdrawal (**Figure 2E**). We observed a similar, albeit milder, effect on AMPK signalling in U2OS cells with siRNA-mediated knockdown of OTULIN, where particularly the levels of ACC and RAPTOR phosphorylation at 0.5-2 hours of starvation were increased (**Figure 2F**). This is consistent with our data from HOIP- and LUBAC-deficient cells and shows that OTULIN is a negative regulator of AMPK signalling.

LUBAC and OTULIN are critical regulators of TNF-induced IKK signalling, NF-κB activation, and cell death^3^, and TNF and NF-κB signalling have been linked to modulation of AMPK signalling^58^. Despite these possible links, we observed no activation of IKKα/β or the related IKK family kinase TBK1 during 6 hours of glucose starvation (**Figure S2C**). While AMPK was phosphorylated and activated as expected during starvation, there was no observable, concomitant phosphorylation of IKKα/β (S176/S180) or TBK1 (S172), nor could any phosphorylation (S32) or degradation of IκBα, the main substrate of IKKα/β, be observed (**Figure S2C**). Addition of TNF-neutralising antibodies to the culture medium during glucose starvation for up to 6 hours also had no bearing on AMPK signalling (**Figure S2D**), whereas it effectively neutralised IκBα phosphorylation and degradation as well as NF-κB p65/RelA phosphorylation (S536) induced by TNF added to the medium. Moreover, we did not observe any sensitisation to starvation-induced cell death upon glucose withdrawal in HOIP-KO cells for up to 24 hours (**Figure S2E**). Together, these data show that neither IKK nor autocrine TNF signalling contribute to the AMPK activation observed upon glucose starvation, hence corroborating that the roles observed for LUBAC and OTULIN in the AMPK pathway are independent of their functions in TNF signalling, IKK activation, and regulation of cell death.

Collectively, our data reveal that LUBAC and OTULIN are *bona fide* regulators of AMPK signalling and that LUBAC’s E3 ligase activity promotes AMPK signalling, while OTULIN restricts it, implying a key role of M1-linked Ub in the regulation of AMPK signalling.

### M1-Linked Ub Signalling is Important for Physiological AMPK Activation and Survival Upon Energetic Stress

The importance of LUBAC and OTULIN in AMPK signalling in cells prompted us to investigate the physiological roles of LUBAC and OTULIN in AMPK signalling. Germline ablation of *Hoip* leads to embryonic lethality in mice^17^. We therefore generated CreERT2-*Hoip*^flox/flox^ mice, in which *Hoip* can be ablated in all cells by tamoxifen administration. Tamoxifen administration led to significant weight loss after 8 days in *Hoip*-deficient mice (**Figure S3A-D**), preventing longer-term studies in these animals. For this reason, we isolated hepatocytes from livers of the *Hoip*-deficient mice (**Figure S3A**) and starved them of glucose *ex vivo.* Consistent with our findings from cell lines, loss of HOIP led to blunted AMPK signalling in primary hepatocytes in response to glucose starvation, observed as reduced phosphorylation of ACC, RAPTOR, AMPKα1/2 and β1, as well as ULK1, compared with controls (**Figure 3A**). Surprisingly, the HOIP-deficient hepatocytes were also less sensitive to AMPK activation by the small-molecule allosteric activator MK-8722^59^ (**Figure 3A**), showing that LUBAC promotes AMPK signalling in primary murine hepatocytes in response to both glucose starvation and direct allosteric activation.

**Figure 3.**
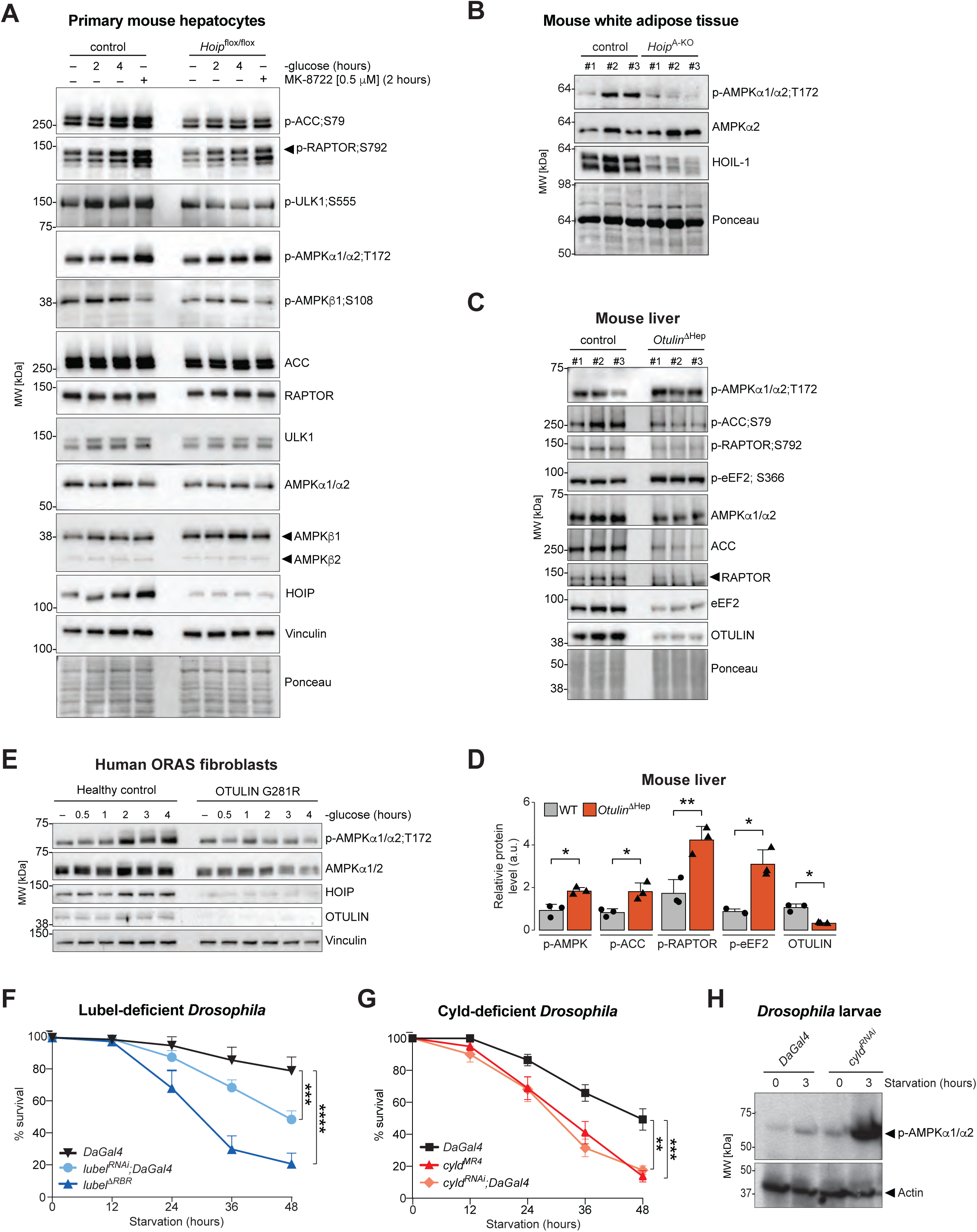
M1-Linked Ub Signalling is Important for Physiological AMPK Activation and Survival Upon Starvation. A) Primary hepatocytes isolated from 10–14-week-old female control or CreERT2- *Hoip*^flox/flox^ mice eight days after initiation of tamoxifen treatment. Cells were cultured overnight in complete medium before undergoing glucose starvation or treatment with the AMPK activator MK-8722 as indicated. Lysates were examined by immunoblotting. B) Lysates from white adipose tissue from control or *Adipoq*Cre-*Hoip*^flox/flox^ mice were analysed by immunoblotting. C) Whole liver lysates from 9-day old control and *Otulin^Δ^*^hep^ mice were analysed by immunoblotting. D) Densitometric analysis of the immunoblot from (C). Bars represent mean signal intensity of phosphoprotein bands normalised to their corresponding total protein bands and ponceau loading ± SEM (n=3). Two-sided Student’s *t*-test was used to determine statistical significance. *p<0.05; **p<0.01. E) Human fibroblasts isolated from an OTULIN-related autoinflammatory syndrome (ORAS) patient with a homozygous OTULIN-G281R^16^ and a healthy control were starved for glucose starvation as indicated. Lysates were analysed by immunoblotting. F) Adult control *DaGal4 Drosophila* flies and mutant *lubel^ΔRBR^* and UAS-*lubel^RNAi^* flies were starved by nutrient withdrawal for up to 48 hours, and their survival was monitored. Data represent mean ± SEM from at least three independent experiments. Two-way ANOVA and the Tukey’s multiple comparison test was used to determine statistical significance. ***p<0.001; ****p<0.0001. G) Adult *DaGal4* control flies, CYLD mutant flies (*cyld^MR4^*), and UAS-*cyld^RNAi^* flies (*cyld^RNAi^*) were starved and analysed as in (F). Data represent mean ± SEM from six independent experiments. Two-way ANOVA and the Tukey’s multiple comparison test was used to determine statistical significance. **p<0.01; ***p<0.001. H) Third instar control *DaGal4* or *cyld^RNA^*^i^ *Drosophila* larvae were starved for three hours and lysates were analysed by immunoblotting. All blots show a single experiment representative of three biologically independent replicates.

To investigate the impact of LUBAC deficiency on physiological AMPK signalling in another metabolic tissue, we used *Hoip*^flox/flox^ mice crossed with *Adipoq*Cre-expressing mice, leading to specific ablation of *Hoip* in adipocytes in which AMPK is a key regulator of metabolism^60^. We analysed AMPK activation in subcutaneous white adipose tissue (WAT) from 22–23-week-old male *Hoip*^flox/flox^-*Adipoq*Cre (*Hoip*^A-KO^) mice and controls that were fed a normal diet. Remarkably, in the fed state with no starvation, we observed a clear overall reduction in AMPK T172 phosphorylation and activation in WAT from *Hoip*^A-KO^ mice (**Figure 3B**). HOIP protein was undetectable in immunoblots of WAT samples from both WT and *Hoip*^A-KO^ mice, but as a proxy for LUBAC loss we observed that HOIL-1 levels were clearly reduced in the *Hoip*^A-KO^ samples (**Figure 3B**), and genotyping confirmed the genotype of the *Hoip*^A-KO^ mice to be *Hoip*^flox/flox^-Cre^+^ (**Figure S3E**), in combination indicating effective ablation of *Hoip*. This demonstrates that loss of LUBAC impairs AMPK activation *in vivo* under physiological conditions.

We continued to investigate if OTULIN also acted as an *in vivo* regulator of AMPK. Mice with hepatocyte-specific deletion of OTULIN (*Otulin*^ΔHep^) display hepatic and systemic metabolic disturbances, including neonatal steatosis, reduced glycogen content, and dysregulation of mTORC1 signalling^28^. In livers from 9-day-old *Otulin*^ΔHep^ mice, we observed clearly elevated AMPK T172 phosphorylation and relatively normal total AMPKα1/2 levels, also in the absence of any imposed energetic stress (**Figure 3C-D**), similar to the conditions in the HOIP-deficient WAT. Of note, for downstream factors of the AMPK signalling pathway, including ACC, RAPTOR, and eE2F, we observed reduced overall protein expression (**Figure 3C**), possibly due to AMPK-mediated inhibition of protein synthesis via eEF2K-eEF2 (**Figure 3C-D**), resembling the observed alterations in the mTORC1 pathway in the *Otulin*^ΔHep^ mice^28^. This indicates that both AMPK and mTORC1 signalling pathways are severely dysregulated in *Otulin*^ΔHep^ livers. Intriguingly, when normalising for the overall reduced protein expression, we observed consistent increases in phosphorylation of the AMPK pathway substrates ACC, RAPTOR, and eEF2 (**Figure 3D**), supporting that OTULIN is a *bona fide* negative regulator of AMPK activation and downstream signalling in cells and *in vivo.* Together, this shows that both LUBAC and OTULIN are physiological regulators of AMPK activation, even in the absence of energetic stress, suggesting they are also important for the physiological steady-state activation and fine-tuning of signalling.

Given the clear impact of LUBAC and OTULIN loss on AMPK activation in mice, we wondered if AMPK signalling was also dysregulated in human disease associated M1-linked Ub dysregulation. We investigated AMPK signalling in primary human skin fibroblasts from an ORAS patient with an OTULIN^G281R^ mutation^16^. These ORAS fibroblast phenocopy LUBAC-deficient cells due to degradation of HOIP concomitant to loss of OTULIN^16^. Strikingly, we observed clearly reduced AMPK activation in the ORAS fibroblasts in response to glucose starvation compared with fibroblasts from a healthy donor (**Figure 3E**). This shows that M1-linked Ub signalling is also required for proper AMPK regulation in primary human cells and that dysregulated AMPK signalling may contribute to the pathogenesis of ORAS and LUBAC deficiency, in which normal M1-linked Ub signalling is disrupted.

Having observed the impact of LUBAC and OTULIN on AMPK activation in primary cells and tissues, we wanted to investigate if M1-linked Ub conjugation and deconjugation was also involved in the physiological response to starvation *in vivo*. For this purpose, we tested the response to starvation in the fruit fly *Drosophila melanogaster*, in which both AMPK^61^ and LUBAC, known as Lubel^62,63^, are conserved. Control flies, flies ubiquitously expressing Lubel-targeting RNAi transcripts under the daughterless-Gal4 (daGal4) driver (*lubel^RNAi^* flies), and flies with genetic deletion of the catalytic RBR ubiquitin ligase domain of Lubel (*lubel*^Δ^*^RBR^* flies) were starved by nutrient withdrawal for up to 48 hours and their survival monitored. Strikingly, the survival rate of *lubel^RNAi^* flies was significantly reduced compared with control flies (**Figure 3F**). The survival rate was further reduced for *lubel*^Δ^*^RBR^* flies with less than 25% of flies surviving 48 hours of starvation (**Figure 3F**). This demonstrates that Lubel – and Lubel’s ubiquitin ligase activity, i.e. M1-linked Ub conjugation – are critically important for the physiological response to starvation. We then tested if deconjugation of M1-linked Ub was also important for the starvation response. While OTULIN is not conserved in flies, the DUB Cyld interacts with Lubel and is the main negative regulator of M1-linked Ub signalling in flies^62,63^. Surprisingly, when we starved Cyld-deficient flies, either RNAi-mediated knockdown flies (*cyld^RNAi^*) or flies with genetic ablation of Cyld (*cyld^MR4^*), we also observed reduced survival rates compared to control flies (**Figure 3G**), similar to Lubel-deficient flies. Starvation of Cyld-deficient fly larvae, where a more synchronised starvation and hence signalling analysis is achievable, showed hyper-phosphorylation of AMPK T172 in the absence of Cyld compared with controls (**Figure 3H**), akin to the loss of OTULIN in cells. This demonstrates that Cyld is as a negative regulator of AMPK activation in *Drosophila* and implies that M1-linked Ub conjugation and deconjugation must be delicately balanced to ensure proper and productive AMPK signalling for survival during starvation in *Drosophila*.

In conclusion, our data show that AMPK activation is controlled by LUBAC and OTULIN in primary mouse hepatocytes, *in vivo* in mice, and in ORAS patient cells. Moreover, we find that Lubel and Cyld are both required for survival during starvation in *Drosophila melanogaster*, showing that M1-linked Ub chains are critical for the protective physiological response to starvation. Collectively, these results highlight the physiological importance of M1-linked Ub signalling in AMPK signalling and the response to starvation across species and in human patients.

### AMPKα and β Subunits Are Putative Substrates of LUBAC Ubiquitination

Having discovered the importance of LUBAC and OTULIN for regulating AMPK signalling, we investigated the mechanism by which they control AMPK. AMPKα has been reported as a substrate of K63-linked ubiquitination, regulating its activity^64^. We therefore tested if AMPK was found in a M1-linked Ub-containing complex and if LUBAC was able to ubiquitinate AMPK subunits. For this purpose, we employed an M1-linked Ub-enrichment strategy using either a new M1-linked Ub-specific binding construct that we devised, called the M1-Trap, which is based on the catalytically inactive OTU domain of OTULIN (OTULIN^80–348;C129A^), or the previously described M1-linkage-specific Ub-binder (M1-SUB) based on NEMO’s UBAN domain^57^. Both domains bind M1-linked Ub chains with high affinity and specificity^6,65^. Of note, the M1-Trap does not contain the OTULIN N-terminus, and hence does not interact directly with AMPK (**Figure S4A**). We co-expressed FLAG-tagged AMPKα1 and α2 in HEK293T cells, together with HOIP, HOIL-1 and OTULIN, and immunoprecipitated the FLAG-tagged AMPKα1/α2 complexes. When co-expressed with HOIP and HOIL-1, FLAG IP of AMPKα1/α2 co-immunoprecipitated a clear smear of M1-linked Ub chains (**Figure 4A**), indicating that LUBAC forms M1-linked Ub chains on one or more proteins found in complex with AMPK. These chains could be completely removed by co-expression of WT OTULIN, whereas they accumulated in the AMPKα1/α2 complexes when co-expressed with catalytically inactive OTULIN^C129A^ (CA), which protects and stabilises M1-linked chains^6,57^ (**Figure 4A**, compare lanes 4 and 5). This shows that the Ub chains co-precipitating with AMPK were indeed M1-linked, and that AMPK can be found in a complex containing M1-linked Ub chains.

**Figure 4.**
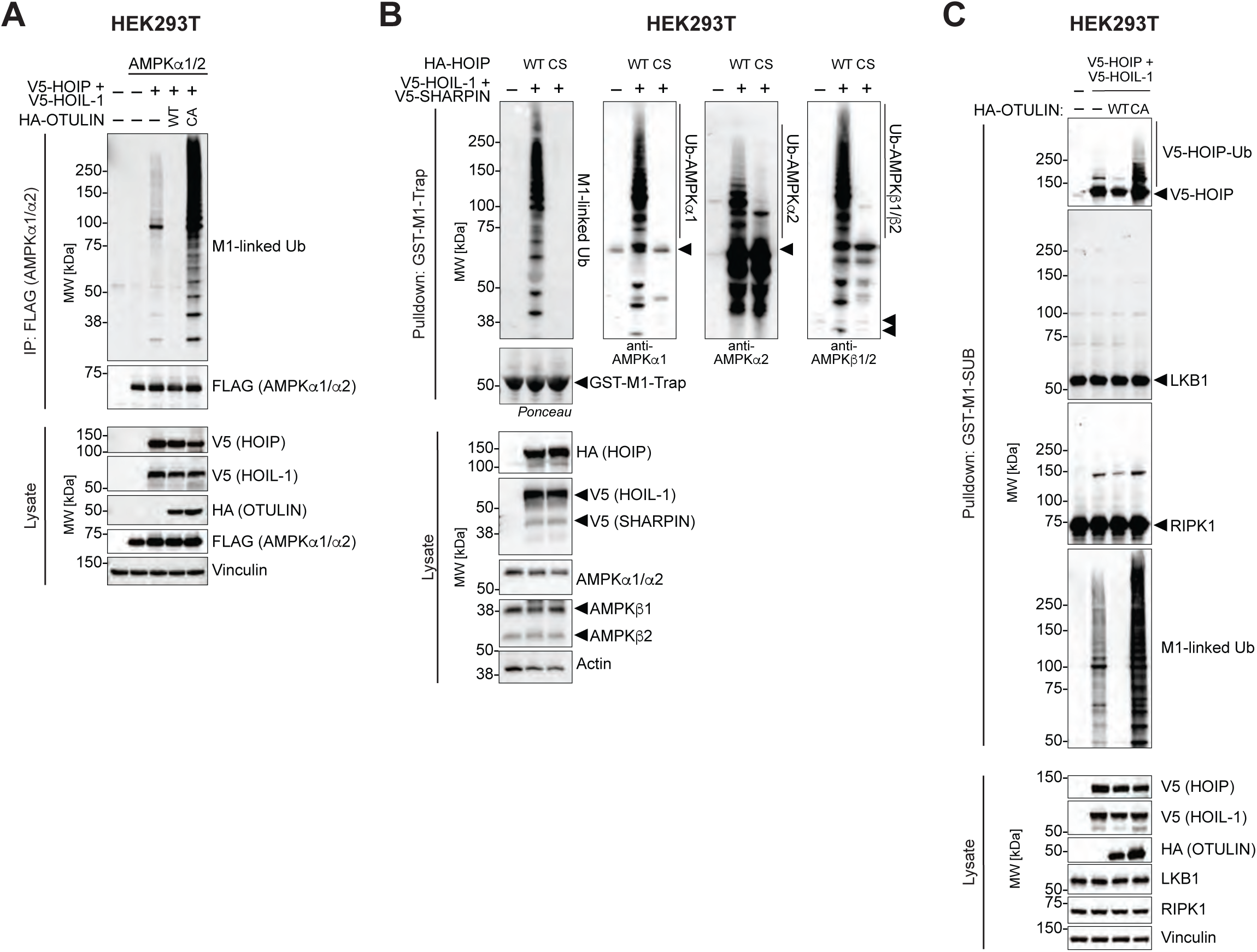
AMPKα and β Subunits Are Putative Substrates of LUBAC Ubiquitination. A) HEK293T cells were transfected with empty vector or FLAG-tagged AMPKα1 and AMPKα2 together with LUBAC (HOIL-1 and HOIP) and either WT OTULIN or catalytically inactive OTULIN-C129A. FLAG-tagged AMPKα1/AMPKα2 was immunoprecipitated, and samples were analysed by immunoblotting. B) HEK293T cells were transfected with empty vector or LUBAC (HOIL-1, SHARPIN, and either WT HOIP or catalytically inactive HOIP-C885S). Endogenous M1-linked Ub chains were affinity-precipitated using recombinant GST-M1-Trap, and the samples were analysed by immunoblotting. C) HEK293T cells were transfected with empty vector or LUBAC (HOIL-1 and HOIP) together with either WT OTULIN or catalytically inactive OTULIN-C129A. Endogenous M1-linked Ub chains were affinity-precipitated using recombinant GST-M1-SUB, and samples were analysed by immunoblotting. All blots show a single experiment representative of three biologically independent replicates, except (C) which is representative of two.

We then tested if LUBAC could ubiquitinate AMPK. Expression of LUBAC with either WT or catalytically inactive HOIP^C879S^, followed by enrichment of M1-linked Ub chains using GST-M1-Trap, revealed that LUBAC effectively ubiquitinated endogenous AMPKα1, α2, and β1/2, which all appeared as smears of high-molecular weight, modified proteins with reduced SDS-PAGE migration on the immunoblots (**Figure 4B**). LUBAC-mediated AMPK ubiquitination was completely dependent on HOIP’s catalytic activity as expression of LUBAC with the HOIP^C879S^ (CS) mutant failed to induce AMPK ubiquitination (**Figure 4B**), indicating that the modified species of AMPK were indeed modified with M1-linked Ub chains in a LUBAC-dependent manner (**Figure 4B**). AMPK ubiquitination was, however, independent of glucose starvation in these experiments (**Figure S4B**). When ectopically expressed, LUBAC also ubiquitinated itself (HOIP autoubiquitination), but was unable to ubiquitinate LKB1, the immediate upstream kinase of AMPK, or RIPK1, a substrate of LUBAC during TNF signalling^13,20^ (**Figures 4C and S4B**), even in the presence of catalytically inactive OTULIN^C129A^ to stabilise M1-linked Ub chains (**Figure 4C**), indicating that LUBAC ubiquitinates AMPKα/β with a degree of selectivity.

Together, these data show that LUBAC can selectively ubiquitinate AMPKα and β subunits, providing a potential mechanism of regulation of AMPK signalling by M1-linked Ub chains.

### LUBAC Controls the Kinase Activity and Activation Threshold of AMPK

Based on our findings that AMPK, LUBAC, and OTULIN form a complex in cells and that LUBAC can conjugate M1-linked Ub chains to AMPK, we further investigated the mechanism by which LUBAC controls AMPK signalling. Ubiquitination of AMPK has mainly been reported to control its protein level or stability^66–68^. However, we did not observe any apparent differences in AMPKα1, α2, β1, β2, or γ1 levels between WT and HOIP-KO U2OS cells at steady-state, nor did we, using a cycloheximide chase, observe any changes in stability or turnover of these AMPK subunits during glucose starvation (**Figure 5A**). We therefore hypothesised that M1-linked Ub chains regulate AMPK signalling in a non-degradative manner.

**Figure 5.**
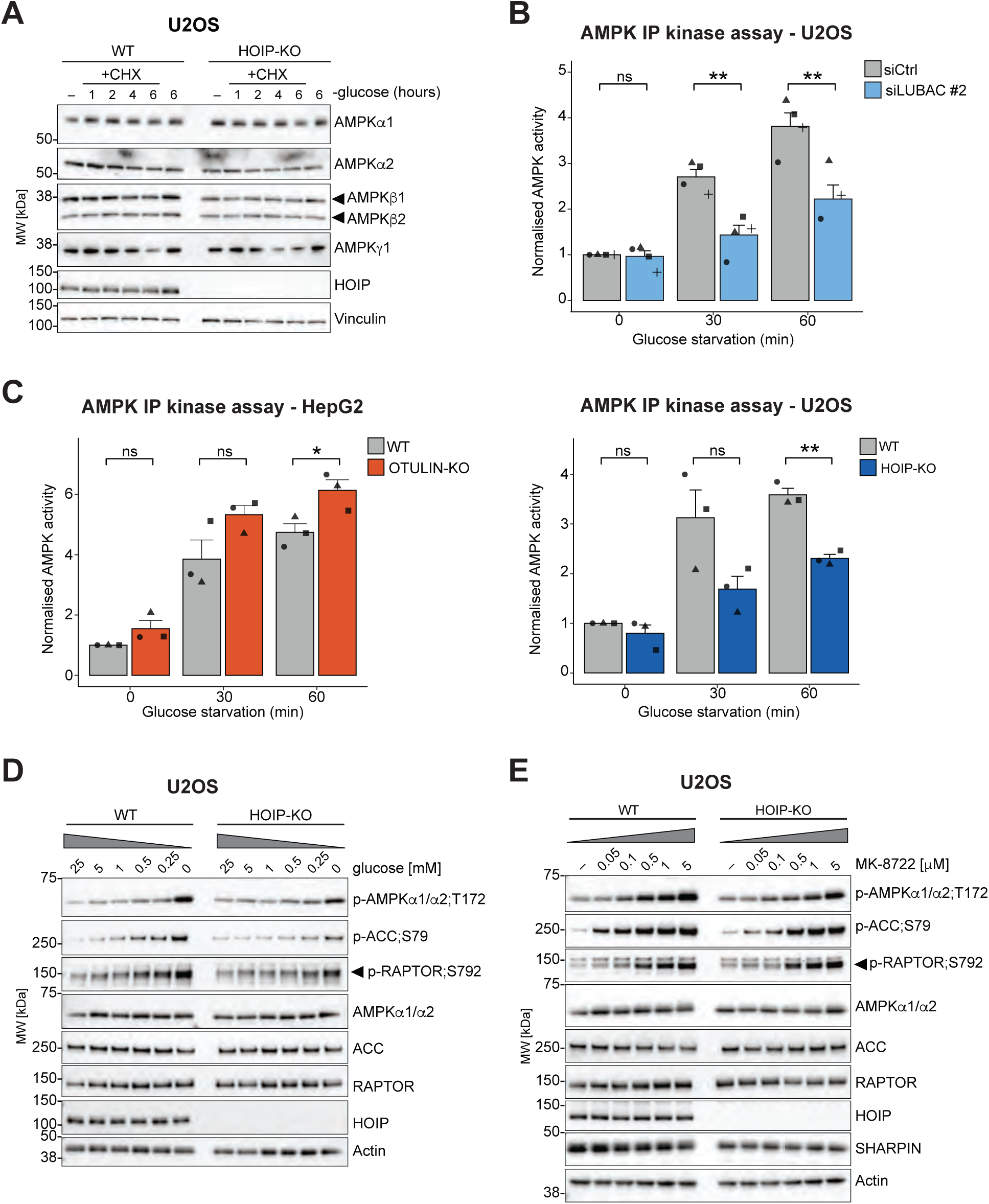
LUBAC Controls the Kinase Activity and Activation Threshold of AMPK. A) U2OS WT and HOIP-KO cells were starved and co-treated with cycloheximide (CHX) to block protein synthesis as indicated. Samples were analysed by immunoblotting. B) AMPKα1/α2 was immunoprecipitated from U2OS cells transfected with either siRNAs targeting HOIP (oligo #2), HOIL-1, and SHARPIN (siLUBAC #2) or non-targeting control (siCtrl) (top) or from U2OS WT or HOIP-KO cells (bottom), after glucose starvation as indicated. AMPK activity was measured by kinase assay with ^33^P-labelled ATP and normalised to untreated control. Bars represent mean activity ± SEM of three biologically independent replicates. Two-sided Student’s *t*-test was used to determine statistical significance. ns, not significant; **p < 0.01. C) HepG2 WT and OTULIN-KO cells were starved for glucose and analysed as in (B). ns, not significant; *p-value < 0.05. D) U2OS WT and HOIP-KO cells were grown with medium containing gradually decreasing concentrations of glucose for 60 minutes and analysed by immunoblotting. E) U2OS WT and HOIP-KO cells were treated with the AMPK activator MK-8722 at increasing concentrations for 60 minutes and analysed by immunoblotting. All blots show a single experiment representative of three biologically independent replicates.

To assess if LUBAC regulated AMPK kinase activity, we performed quantitative IP-coupled kinase assays, in which AMPK was immunoprecipitated from cells and its intrinsic activation status quantified by a kinase assay of the purified kinase with a synthetic substrate peptide and radiolabelled ATP^69^. Intriguingly, the activity of the AMPK that immunoprecipitated from U2OS cells with siRNA-mediated knockdown of LUBAC or genetic ablation of HOIP after glucose starvation was significantly reduced compared with the respective controls (**Figure 5B**). Conversely, the activity of the AMPK immunoprecipitated from OTULIN-KO HepG2 cells after glucose starvation was significantly increased (**Figure 5C**). These data show that LUBAC and OTULIN regulate AMPK’s kinase activity, consistent with the effects on AMPK signalling we observed in our immunoblot analyses (**Figure 2**).

We noticed that the effect of LUBAC deficiency on AMPK activity was evident already at 30 minutes of starvation, so we asked whether the ability of AMPK to sense immediate or small reductions in glucose or energy levels was affected in LUBAC-deficient cells. To investigate this, we subjected U2OS WT and HOIP-KO cells to medium containing decreasing amounts of glucose (from 25 mM to 0 mM) and found that loss of HOIP increased the activation threshold of AMPK noticeably. In WT cells, we observed AMPK activation, evident as increased AMPK T172, ACC S79, and RAPTOR S792 phosphorylation, starting at glucose concentrations of ∼5-1 mM (**Figure 5D**). In contrast, we only observed phosphorylation exceeding background of AMPK, ACC, and RAPTOR starting at glucose concentrations of ∼0.5-0.25 mM in HOIP-KO cells (**Figure 5D**). A similar increase in activation threshold was observed in U2OS cells depleted for LUBAC with siRNAs (**Figure S5A**). This shows that lower glucose levels (∼0.25 mM) are required for AMPK activation in LUBAC-deficient cells compared with WT cells (∼1 mM), indicating that LUBAC controls the activation threshold of AMPK in response to glucose starvation.

Canonically, AMPK is activated by elevations in intracellular AMP:ATP or ADP:ATP ratios, but AMPK can also be activated after glucose starvation via direct glucose-sensing without changes in AMP:ATP or ADP:ATP ratios^70^. We therefore measured adenine nucleotide levels during glucose starvation using LC-MS/MS and calculated the adenylate energy charge (AEC), which is an index of intracellular energy status^71^, to determine if loss of either HOIP or OTULIN affected the AMP:ADP:ATP ratios (**Figure S5B-C**). Energetic stress, i.e. an increase in AMP:ATP and ADP:ATP ratios, leads to a decrease in AEC. In both WT U2OS and WT HepG2 cells, the AEC decreased significantly over 30-60 minutes of glucose starvation (**Figure S5B-C**), showing that these cells experience a drop in energy levels and increased AMP/ADP:ATP ratios upon glucose starvation. Importantly, glucose withdrawal from HOIP- or OTULIN-KO cells was accompanied by similar decreases in AEC (**Figure S5B-C**), demonstrating that HOIP- and OTULIN-deficient cells experience a similar drop in energy levels and increases in relative AMP and ADP levels as WT cells, and hence implying that LUBAC and OTULIN regulate canonical AMPK activation. This suggests that the blunted AMPK activation and increased activation threshold in HOIP-deficient cells is not caused by lower AMP or ADP levels in these cells (**Figure S5B**). Instead, it indicates that the ability of AMPK to become activated in response to elevated AMP levels is impaired in the absence of LUBAC.

Exploring the LUBAC-mediated AMPK activation mechanism further, we tested if LUBAC also affected AMPK activation independently of changes in nucleotide levels. In primary hepatocytes, we already observed that *Hoip*-ablated cells were less sensitive to the small-molecule allosteric AMPK activator MK-8722^74^. MK-8722 binds to the ADaM site between the kinase domain in the α-subunit and carbohydrate binding module in the β-subunit and protects the phosphorylated T172 against phosphatases in a nucleotide-independent manner^72,73^. We tested MK-8722 potency in HOIP-KO U2OS cells and found that loss of HOIP led to impaired responsiveness of AMPK to MK-8722 (**Figure 5E**), similar to our observations for glucose starvation. MK-8722 treatment caused AMPK, ACC, and RAPTOR phosphorylation starting at a concentration of 0.05-0.1 μM in WT cells, whereas 0.1-0.5 μM was required for a similar response in HOIP-KO cells (**Figure 5E**). This demonstrates that LUBAC is required for proper AMPK activation in response to both energetic stress and direct activation by allosteric activators.

Collectively, we find that LUBAC and OTULIN control the kinase activity of AMPK but do not affect the cellular adenine nucleotide ratios in response to glucose starvation, indicating that M1-linked Ub-dependent regulation of AMPK operates downstream of changes in energy charge. Here, LUBAC controls not only the amplitude of AMPK activation but also the threshold of activation in response to both glucose starvation and direct activation by MK-8722.

### Phosphoproteomic Analysis Reveals the LUBAC-Dependent Signalling Outcomes After Glucose Starvation

Energetic stress activates a range of adaptive processes in cells, including AMPK signalling, which regulates various cellular responses. To investigate the role of M1-linked Ub signalling in the broader context of starvation response, we conducted a mass spectrometry-based phosphoproteomic analysis of glucose-starved U2OS WT and HOIP-KO cells (0, 10, 30, 60, 120 minutes of starvation) using tandem mass tag (TMT) labelling and Fe(III)-based phospho-peptide enrichment (**Figure 6A**). We identified more than 20,000 phosphorylation sites (phosphosites) distributed across more than 4,500 proteins (**Figure 6B**). Glucose starvation resulted in significant changes to the phosphorylation status of hundreds of sites (**Figure S6A-B and Table S3**), but the overall phosphoproteomic landscape between WT and HOIP-KO cells remained relatively comparable even in response to glucose withdrawal (**Figure S6B-C**). This indicates that the absence of HOIP perturbs specific cellular signalling events during starvation rather than causing a widespread impairment of signalling pathways. We applied fuzzy c-means (FCM) clustering^74^ of the WT samples to group phosphosites into four clusters termed “*Fast up*”, “*Slow up*”, “*Fast down*”, and “*Slow down*” based on their dynamics (**Figure 6C**). The autophosphorylation sites of AMPKβ1 (PRKAB1; S108) and AMPKβ2 (PRKAB2; S108), as well as the AMPK sites on ACC (ACACA; S80) and RAPTOR (RPTOR; S722), were part of the “*Slow up*” cluster (**Figure 6C**; left). Functional enrichment analysis of KEGG pathways across all clusters confirmed that AMPK signaling was significantly enriched in the “*Slow up*” cluster (**Figure 6C**; right, **Table S4**). Subsequently, we analysed the phosphosites in the “*Slow up*” cluster using the Kinase-Substrate Enrichment Analysis (KSEA) tool^75^, which infers kinase activities based on fold-changes of phosphosites compared with control. KSEA revealed that AMPKα1 (PRKAA1) was the most active kinase in the “*Slow up*” cluster, significantly enriched in both WT and HOIP-KO cells along with 9 other kinases that we could assign significant positive activity scores to, including glycogen synthase kinase 3 β (GSK3β; GSK3B), ribosomal S6 kinase A1 (S6K1; RSP6KA1), and Ca^2+^/Calmodulin-dependent protein kinase 2 alpha (CamK2α; CAMK2A) (**Figures 6D and S6D**). Importantly, the normalised kinase activity (reflected in the z-score) for AMPKα1 (PRKAA1) was consistently lower in HOIP-KO compared with WT, demonstrating an impaired and delayed AMPK response in HOIP-KO cells upon starvation (**Figures 6D and S6D**), in agreement with our other data in this study. Interestingly, the mTOR kinase was more active in the HOIP-KO during glucose starvation within this cluster, demonstrating significant activation at 10 minutes (**Figures 6D and S6D**). Given that AMPK inhibits mTOR activity through TSC2 and RAPTOR phosphorylation, this observation is consistent with the lower activation of AMPK observed in HOIP-KO cells, resulting in reduced AMPK-mediated inhibition of mTOR. Consistently, S6K1 and PRKCA, which are targets of mTOR, were also more active in HOIP-KO cells than in WT (**Figures 6D and S6D**). Together, this indicates a disruption of kinase kinetics downstream of AMPK in HOIP-KO cells during glucose starvation. Notably, the second most activated kinase in the “*Slow up*” cluster, GSK3β, which also has important roles in modulating glucose metabolism and energy homeostasis^76^, was less influenced by loss of HOIP than AMPK (**Figure 6D and S6D**), indicating that LUBAC primarily and specifically regulates AMPK activation during starvation.

**Figure 6.**
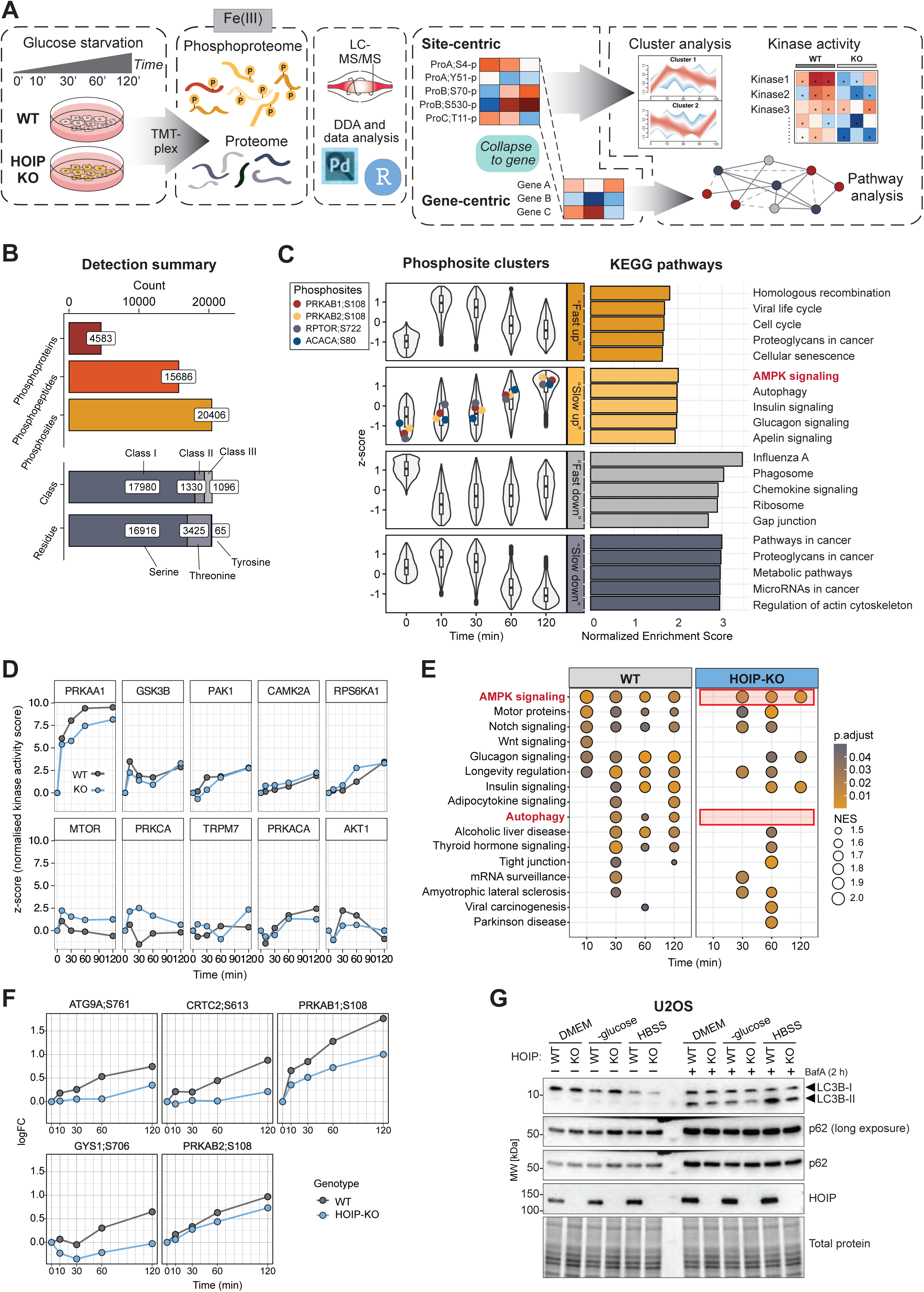
Phosphoproteomic Analysis Reveals the LUBAC-Dependent Signalling Network After Glucose Starvation. A) Workflow schematic of LC-MS/MS-based phosphoproteomics analysis of U2OS WT and HOIP-KO cells glucose-starved for 0-120 minutes (n = 3 per time point). Data were processed with site-centric (for phosphorylation dynamics and KSEA) and gene-centric (for pathway analysis) approaches. B) Top: Bar plots showing total numbers of phosphoproteins, phosphopeptides, and phosphorylation sites (phosphosites) for the entire dataset. Middle: Numbers of Class I, II, and III phosphosites with localization probabilities of >75%, 50-75%, and <50%, respectively. Bottom: Total numbers of phosphosites on serine, threonine, and tyrosine residues. C) Left: Violin plots of four fuzzy c-means cluster profiles of phosphosite z-scores over time in WT cells. Clusters are named by their phosphorylation dynamics: “Fast up”, “Slow up”, “Fast down”, and “Slow down”. Phosphorylated AMPKβ1/β2;S108, RAPTOR;S722, and ACC1;S80 are in the “Slow up” cluster. Right: Bar plots of the top five significantly upregulated KEGG pathway terms in each cluster, identified via Gene Set Enrichment Analysis (GSEA, p < 0.005), with “AMPK signalling” highlighted in the “Slow up” cluster. D) Plots showing kinase activity scores (z-scores) of the top 10 most active kinases in the “*Slow up*” cluster in WT (grey) and HOIP-KO cells (blue) during glucose starvation. Kinase activity scores were determined using the KSEA tool. E) Dot plot visualising which KEGG pathways are significantly enriched in WT or HOIP-KO cells during glucose starvation, analysed by GSEA (p<0.05). Log2 fold-change of phosphosites over untreated cells for each genotype was analysed. The analysis was performed using GSEA on KEGG pathways with normalised enrichment scores (NES) shown by dot size, Dot colour represents the statistical significance (BH-adjusted p-value). F) Log2 fold-changes (LogFC) over untreated cells of phosphosites in ATG9A, CRTC2, AMPKβ1/β2, and GYS1 in WT (grey) and HOIP-KO cells (blue) are plotted for each time point. G) U2OS WT and HOIP-KO cells were starved in glucose-free medium or in Hank’s Balanced Salt Solution (HBSS) and treated with bafilomycin A1 or solvent control for two hours. Lysates were analysed by immunoblotting. Blots show a single experiment representative of three biologically independent replicates.

To direct our analysis of the cellular functions controlled by LUBAC during starvation, we collapsed the phosphosites from the “*Slow up*” cluster into their corresponding genes to assess signalling outcomes at the gene level rather than at the individual phosphosite level (**Figure 6A**). We then explored the differentially regulated pathways between WT and HOIP-KO cells following starvation using KEGG pathway analysis (**Figure 6E and Table S5**). The immediate response to starvation was characterised by pathways related to AMPK, glucagon signalling, and longevity pathways, as well as Wnt signalling and motor protein regulation. The strongest and most immediate response in WT cells after 10 minutes of starvation was AMPK signalling, which was reduced in HOIP-KO cells (**Figure 6E and Table S5**), consistent with our KSEA analysis. HOIP-deficiency also impaired other pathways in the immediate response, including Wnt signalling, consistent with reported functions of M1-linked Ub in the Wnt pathway^7,77^. In the second wave of signalling, 30 minutes of starvation was primarily associated with insulin signalling, autophagy, and longevity pathways (**Figure 6E and Table S5**). Strikingly, we noticed a severe impairment of autophagy signalling in the HOIP-KO cells in response to starvation (**Figure 6E**), consistent with a reduction in activation of AMPK, a key regulator of autophagy^41^, in these cells. Closer inspection of phosphosites associated with autophagy and AMPK signalling in this cluster, revealed several significantly regulated sites that were reduced in HOIP-KO cells, including Autophagy-related protein 9A (ATG9A; S761), CREB-regulated transcription coactivator 2 (CRTC2; S613), AMPKβ1 and β2 (PRKAB1/2; S108), and glycogen synthase 1 (GYS1; S706) (**Figure 6F**). This indicated that the phosphorylation- and AMPK-dependent autophagic response to glucose starvation was impaired in the absence of HOIP. We validated this experimentally by assessing the autophagic response in WT and HOIP-KO cells. Consistent with our phosphoproteomics, we observed an impaired autophagic response to glucose starvation or complete starvation in HBSS-buffer in HOIP-KO cells (**Figure 6G**) To assess autophagic flux, LC3B-I to LC3B-II conversion and p62 degradation were measured in presence or absence of Bafilomycin A1 (Baf A1), which blocks lysosomal acidification. The glucose and HBSS starvation-induced p62 degradation was partially abrogated in HOIP KO cells (without Baf A1 treatment), while the BafA1-mediated accumulation of LC3B-II and p62 was reduced in HOIP-KO cells relative to WT, indicating defective autophagic flux (**Figure 6G**). This points to impaired phosphorylation-dependent induction of autophagy in LUBAC-deficient cells.

Collectively, our phosphoproteomic analysis shows that LUBAC controls a select and specific set of kinase signalling events, primarily AMPK signalling, during the starvation response and identifies LUBAC as a regulator of starvation-induced autophagy, potentially influencing the ability of cells to cope with energetic stress.

### LUBAC Regulates Bioenergetic Metabolism in Response to Starvation

Our observation that LUBAC is required for proper AMPK activation and the autophagic response to starvation implied that LUBAC would also be functionally involved in controlling cellular metabolism and energetics. To test this, we measured bioenergetics in WT and HOIP-KO U2OS cells in complete and glucose-free medium on a Seahorse XF analyser, quantifying oxygen consumption rate (OCR), extracellular acidification rate (ECAR), and derived metrics of mitochondrial aerobic or glycolytic function over time (**Figure S7A-B**). In complete medium, the OCR of HOIP-KO cells was relatively normal and comparable to WT cells, also in response to different perturbations (**Figures 7A and S7A)**. We did observe a slight decrease in maximal respiration in the HOIP-KO cells (**Figure 7A-B and S7A**), potentially suggesting a mild functional impairment of mitochondria in LUBAC-deficient cells. However, we did not observe any difference in ATP-linked respiration between WT and HOIP-KO cells in complete medium (**Figure 7A-B**), indicating that LUBAC does not modulate the proportion of mitochondrial respiration designated to ATP production at steady-state. Overall, this showed no overt defects in bioenergetic metabolism in HOIP-deficient cells in complete medium.

**Figure 7.**
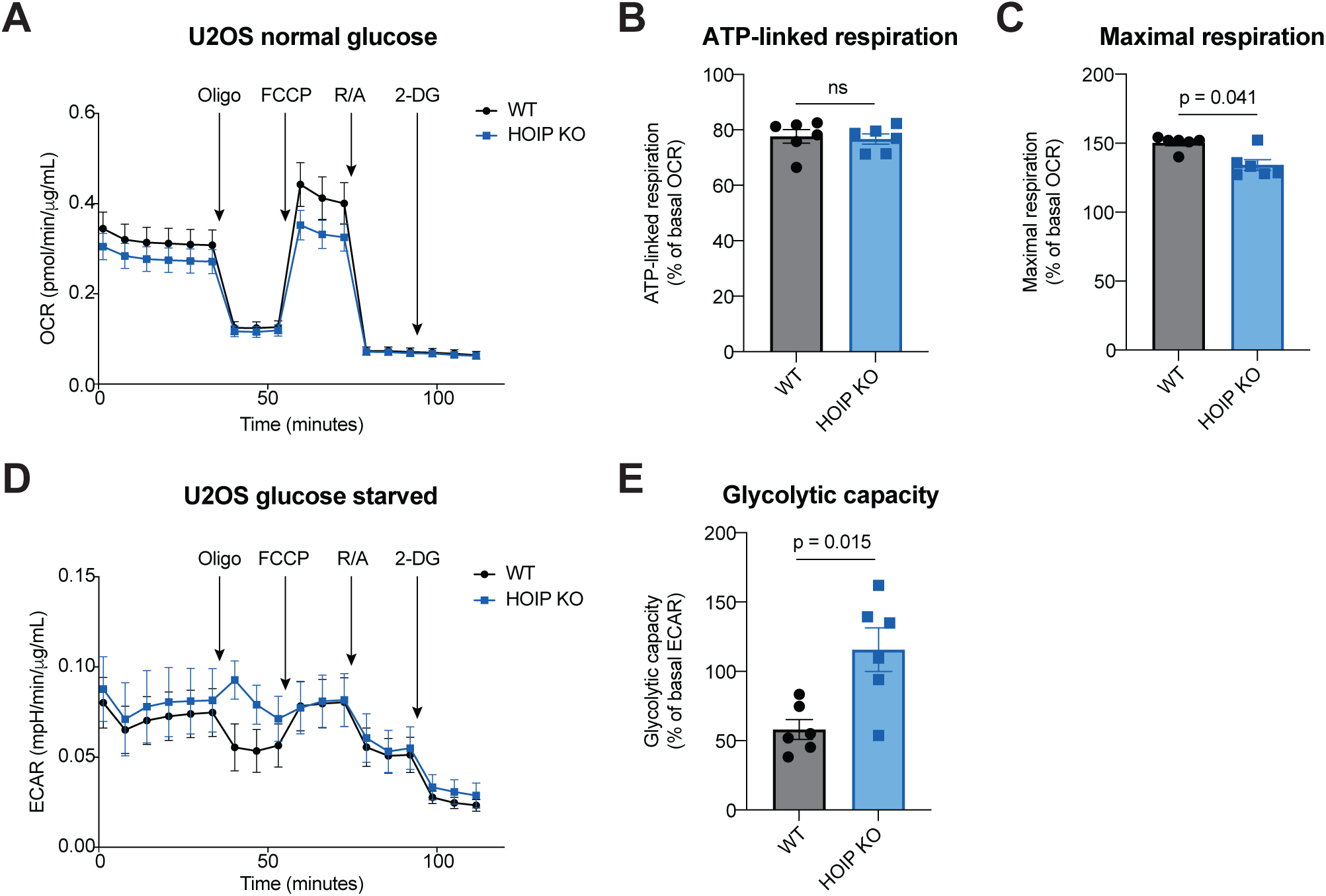
LUBAC Regulates Bioenergetic Metabolism in Response to Starvation. A) Oxygen consumption rate (OCR) in U2OS WT (grey) or HOIP-KO cells (blue) was quantified over time in complete medium with the indicated perturbations. B) Bar plot of ATP-linked respiration in WT and HOIP-KO cells, shown as a percentage of basal OCR following oligomycin injection. C) Bar plot of maximal respiration in WT and HOIP-KO cells, shown as a percentage of basal OCR following FCCP injection. D) Extracellular acidification rate (ECAR) in U2OS WT (grey) or HOIP-KO cells (blue) was quantified over time in glucose-free medium with the indicated perturbations. E) Glycolytic capacity in WT and HOIP-KO cells in glucose-free medium, shown as a percentage of the basal ECAR. Bars represent mean across six independent experiments ± SEM. Statistical testing was performed using a two-tailed non-parametric Mann-Whitney test, with exact p-values indicated.

We then tested how HOIP-KO cells responded to energetic stress. We shifted the cells to glucose-free medium and monitored the ECAR (**Figure 7D and S7B**).

Interestingly, we observed a significant difference in glycolytic capacity between WT and HOIP-KO cells (**Figure 7D-E and S7B**). In glucose-deprived medium, the production of ATP is stunted, which activates AMPK to stimulate increased glycolysis^78,79^. For normal, AMPK-proficient cells in glucose-free medium, glycolysis should operate at nearly maximum capacity, and the cells should therefore not be able to increase their glycolysis markedly when forced to do so by inhibition of respiration by the ATP synthase inhibitor oligomycin. AMPK-proficient cells should therefore have a glycolytic capacity ≤100% (**Figure S7B**). As observed in glucose-starved WT cells, upon oligomycin treatment, there was a drop in ECAR, resulting in a glycolytic capacity of ∼55%, showing that they cannot increase their rate of glycolysis further (**Figure 7D-E**). In contrast, in HOIP-KO cells, glycolytic capacity was increased to ∼115% upon ATP-synthase inhibition, indicating the cells are operating at sub-maximal glycolytic capacity in glucose-free medium (**Figure 7D-E**). This shows that HOIP-KO cells do not switch fully to glycolysis within ∼30-40 minutes of starvation, consistent with a defect in AMPK activation.

In conclusion, our data show that LUBAC is important for controlling not only AMPK signalling in response to starvation, but also the switch to glycolysis and hence the bioenergetic metabolism during energetic stress.

## Discussion

Here, we show that LUBAC is an important regulator of AMPK signalling and the response to starvation, identifying the first metabolic signalling pathway regulated by M1-linked Ub. Our work thus open new avenues of research into the molecular regulation of metabolism by M1-ubiquitination. We demonstrate that LUBAC’s catalytic activity promotes AMPK activation, while OTULIN restricts it, and that both formation and disassembly of M1-linked Ub chains are important for control of AMPK pathway signalling. We reveal that LUBAC and OTULIN interact with heterotrimeric AMPK in cells, and that LUBAC can M1-ubiquitinate AMPKα1/2 and β1/2 subunits. Phosphoproteomics indicate that LUBAC regulates a select set of kinase signalling events during starvation, primarily AMPK signalling and autophagy, suggesting that it is a specific regulator of AMPK. Consistent with our AMPK signalling data, we found that LUBAC controls cellular bioenergetics and is critical for the physiological response to energetic stress in flies. Surprisingly, both Lubel- and Cyld-deficient flies have significantly reduced survival rates in response to starvation, indicating a more complex role for M1-linked Ub in the physiological starvation stress response where assembly and disassembly must be delicately balanced for a productive survival response.

LUBAC and OTULIN are important regulators of IKK activation and cell death in TNF signalling. Interestingly, our data shows that the role of LUBAC and M1-linked Ub in AMPK signalling is not related to TNF, IKK or NF-κB activation, or cell death. We were unable to detect IKK activation during glucose starvation, and blocking TNF signalling also had no impact on AMPK signalling. In addition, we observed no sensitisation to starvation-induced cell death in HOIP-deficient cells for up to 18-24 hours. Collectively, these data indicate that LUBAC (and OTULIN) regulate AMPK activation and signalling directly and independently of their functions in TNF signalling, IKK activation, and cell death.

Loss-of-function mutations in LUBAC or OTULIN cause the severe TNF-driven autoinflammation syndromes LUBAC deficiency and ORAS. These syndromes involve unexplained manifestations linked to glycogen and lipid metabolism, including glycogen storage disease (amylopectinosis) in skeletal and cardiac muscle, lipodystrophy, subcutaneous fat inflammation, and liver steatosis^16,19,23,24,26–28^. Our finding that the M1-linked Ub machinery regulates AMPK signalling, and that loss of either HOIP or OTULIN causes AMPK dysregulation in cells and animal models,

suggests that dysregulated AMPK signalling may contribute to the pathogenesis and clinical manifestations of LUBAC deficiency and ORAS. Consistent with this notion, we observed strongly dysregulated AMPK signalling in fibroblasts from an ORAS patient, clearly implying that AMPK activation is altered in patients. AMPK is a central regulator of both glycogen and lipid metabolism in liver, muscle, and adipose tissue^30,60^, consistent with the idea that dysregulated AMPK signalling could be contributing to the metabolic manifestations in LUBAC deficiency and ORAS. Intriguingly, amylopectinosis observed in LUBAC deficiency patients with HOIP or HOIL-1 mutations^23,24,27^ is phenocopied in *Hoil-1*-deficient mice^80,81^. Recently, downregulation of GYS1 was shown to prevent amylopectinosis in *Hoil-1*-deficient mice^82^. GYS1 is directly phosphorylated and inhibited by AMPK^83^, thereby limiting glycogen synthesis. We find that AMPK activity is reduced in LUBAC deficient cells and tissues, suggesting that AMPK-mediated GYS1 phosphorylation would be impaired, leaving GYS1 more active to promote glycogen synthesis. Consistently, we find that GYS1 phosphorylation is reduced in HOIP-KO cells, although the site identified in our phosphoproteome is not an annotated AMPK site. However, we speculate that reduced LUBAC-mediated AMPK activity could cause GYS1 hyperactivation, leading to aberrant glycogen synthesis and resulting in amylopectinosis in LUBAC deficiency patients and mouse models. Intriguingly, the glycogen content is severely reduced in OTULIN-deficient mouse livers^28^, in which we here show elevated AMPK activity. This supports the notion of AMPK-mediated GYS1 regulation as the explanation for glycogen accumulation in LUBAC deficiency and glycogen depletion in OTULIN deficiency.

Our data demonstrates that LUBAC’s catalytic ubiquitin ligase activity is required for full AMPK activation, meaning that M1-linked Ub chains, possibly conjugated to AMPKα/β subunits, promote AMPK activation and signalling. This suggests that AMPK could be a Ub-activated kinase, akin to IKK. LUBAC-assembled M1-Ub chains promote the activation of IKK facilitating the recruitment of IKK to M1-linked Ub chains via the UBAN domain in the regulatory subunit NEMO^65^. The interaction between NEMO and M1-linked Ub is required for IKK activation, and Ub-binding likely induces a conformational change in IKK, which enables IKK phosphorylation and activation by the upstream kinase TAK1^84,85^. Our data indicate that M1-linked Ub controls the kinase activity and activation threshold of AMPK in response to both glucose starvation (downstream of changes in energy charge) and direct allosteric activation by MK-8722. The observation that M1-linked Ub controls

AMPK activation in response to both increases in AMP/ADP:ATP ratios and allosteric activation, suggests LUBAC regulates a mechanism of AMPK activation involved in multiple modes of activation. We speculate that the M1-linked Ub chains, possibly conjugated to AMPK, could promote AMPK activation by mediating the interaction with or activation by an upstream kinase, e.g. LKB1, by protecting the AMPKα T172 phosphorylation from phosphatases, or by controlling AMPK’s subcellular localisation or complex composition, which would allow regulation of multiple modes of AMPK activation. It will be important to dissect the detailed molecular mechanisms by which M1-linked Ub regulate AMPK activation in future studies.

In addition to regulating AMPK activation, we find that LUBAC also controls cellular bioenergetics in response to starvation. The starvation-induced shift from oxidative phosphorylation to glycolysis was impaired in HOIP-KO cells, consistent with an impaired AMPK-mediated activation of glycolysis in these cells. These data are consistent with findings from HOIP-deficient tumours in mice, which also exhibited lower levels of glycolysis^86^, supporting our notion that LUBAC regulates bioenergetic metabolism via AMPK. The detailed regulation of metabolism by M1-linked Ub, e.g. by metabolomic analysis, will be interesting to dissect in future studies.

In conclusion, our findings conceptually expand the cellular and physiological role of LUBAC and M1-linked Ub to include regulation of metabolic signalling and metabolism, and they reveal atypical ubiquitination as a new level of posttranslational regulation of AMPK.

## Acknowledgements

We would like to thank the DTU Proteomics Core, and in particular facility manager Marie Vestergaard Lukassen and Valdemaras Petrosius for assistance with experimental design, sample preparation, and proteomic analyses, as well as laboratory technician Victoria Nicolaysen for technical assistance. We thank Dr Abdel Atrih at the FingerPrints Proteomics Facility at University of Dundee for nucleotide analyses and the Åbo Akademi University Fly Unit supported by Biocenter Finland for assistance with fly experiments. We also extend our gratitude to Professor David Komander (WEHI, Melbourne, Australia) and Programme Leader Andrew N.J. McKenzie (MRC-LMB, Cambridge, UK) for sharing samples from OTULIN-deficient mice, and Professor Sir Philip Cohen (University of Dundee, UK) for sharing HOIP knock-in MEFs. We are also grateful to Professor Ulrich auf dem Keller (in memoriam) for his support and guidance. This work was supported by a Hallas-Møller Emerging Investigator grant (NNF19OC0054248) from Novo Nordisk Foundation (R.B.D.) an EliteForsk Award (2083-00007B) from The Ministry of Higher Education and Science in Denmark (C.R.E.), the Swiss National Science Foundation (IZSTZ0_223324) (J.D.), an Investigator Award (204766/Z/16/Z) from the Wellcome Trust UK (S.A.H. and D.G.H), the InFLAMES Flagship Programme of the Academy of Finland (#337531) and the Swedish Cultural Foundation (#199483) (A.M), UKRI-Medical Research Council (grant MR/Y014987/1) (B.R.), and the Deutsche Forschungsgemeinschaft (SFB1403 #414786233) (N.P.). The Novo Nordisk Foundation Center for Basic Metabolic Research based at the University of Copenhagen is an independent Research Center and partially funded by an unconditional donation from the Novo Nordisk Foundation (Grant number NNF18CC0034900 and NNF23SA0084103).

## Author Contributions

**CRE:** Conceptualization; Investigation, Formal Analysis; Methodology; Visualization; Data Curation; Funding Acquisition; Writing – Original Draft; Writing – Review and Editing. **ASJ:** Investigation; Methodology; Writing – Review and Editing. **JC:** Investigation; Methodology; Writing – Review and Editing. **AMD:** Investigation; Writing – Review and Editing. **ALA:** Investigation; Writing – Review and Editing. **JR:** Investigation; Methodology; Writing – Review and Editing. **SAH:** Supervision; Writing – Review and Editing. **SNJF:** Investigation; Writing – Review and Editing. **MS:** Investigation; Methodology; Data Curation; Writing – Review and Editing. **SPK:** Investigation; Writing – Review and Editing. **XH:** Investigation; Writing – Review and Editing. **AEF:** Validation; Writing – Review and Editing. **KN:** Investigation; Writing – Review and Editing. **DP:** Investigation; Writing – Review and Editing. **JK:** Resources; Writing – Review and Editing. **MD:** Investigation; Writing – Review and Editing. **JG:** Investigation; Writing – Review and Editing. **LV:** Resources; Writing – Review and Editing. **GvL:** Resources; Supervision; Writing – Review and Editing. **NP:** Resources; Supervision; Writing – Review and Editing. **LBF:** Resources; Supervision. Writing – Review and Editing. **MGH:** Resources, Methodology; Supervision; Writing – Review and Editing. **JD:** Resources; Methodology; Supervision; Writing – Review and Editing. **BJR:** Resources; Supervision; Writing – Review and Editing. **DGH:** Resources; Supervision, Writing – Review and Editing. **AM:** Resources; Methodology; Investigation; Supervision; Writing – Review and Editing. **KS:** Resources; Methodology; Supervision; Writing – Review and Editing. **RBD:** Conceptualization; Formal Analysis; Methodology; Visualization; Supervision; Funding Acquisition; Project Administration; Writing – Original Draft; Writing – Review and Editing.

---

**Supplementary Figure S1. Relating to Figure 1.**
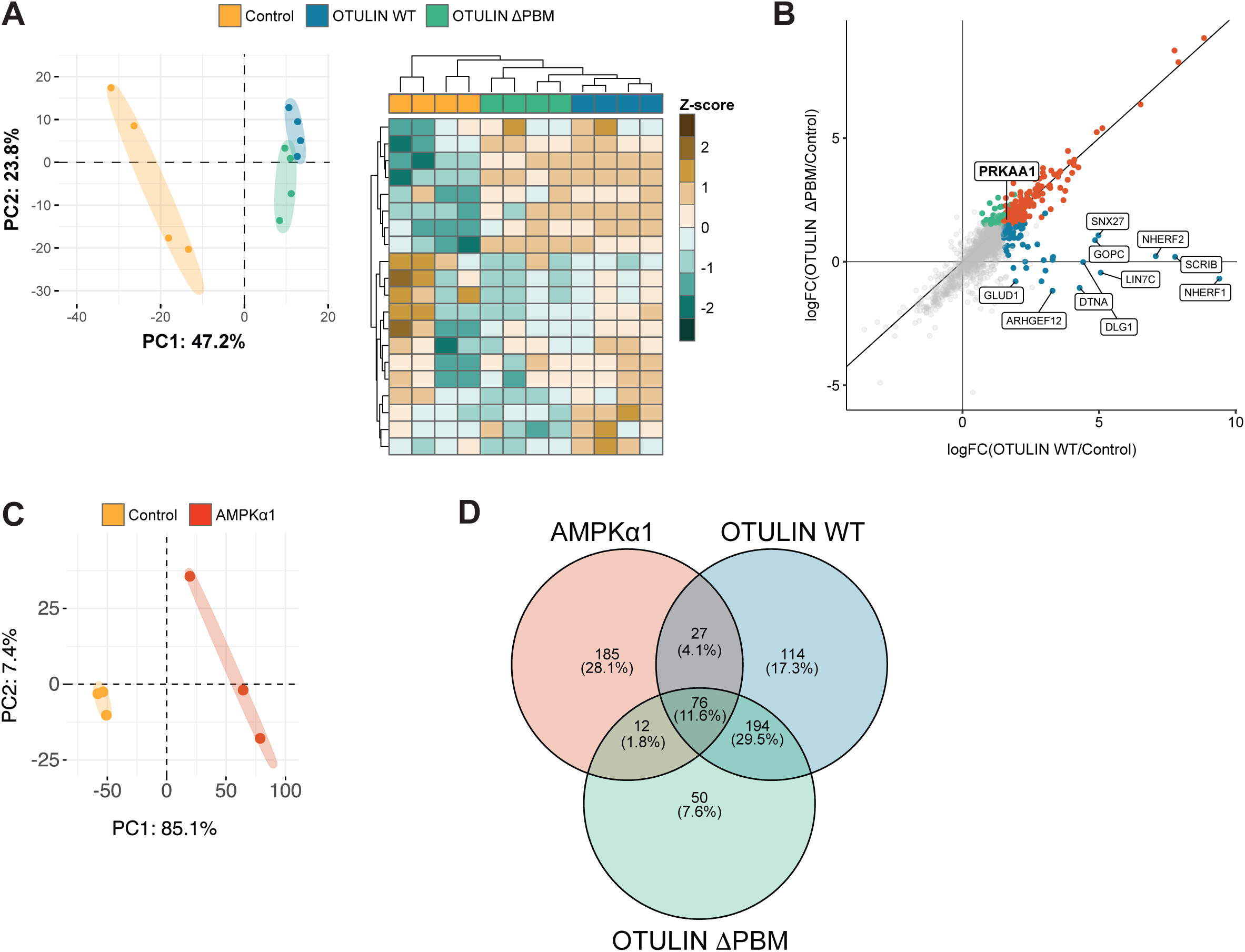
A) Left: Principal Component Analysis (PCA) of mass-spec analysed pulldowns from HEK293T cells overexpressing empty vector (Control) or 2xHA-2xStrep-tagged OTULIN WT or OTULIN ΔPBM. The plot shows PC1 and PC2 explaining 71% of the total variance. Right: Heatmap of normalised protein intensities with hierarchical Ward’s D2 linkage clustering of samples based on Jaccard distances. The heatmap is built using k-means clustering of proteins into 20 clusters. B) Biplot of log2 fold-change (logFC) of proteins in OTULIN ΔPBM over Control versus OTULIN WT over Control. Proteins most significantly enriched in OTULIN WT but not OTULIN ΔPBM are labelled, as well as AMPKα1, which is significantly enriched in both pulldowns. A BH-adjusted FDR<0.05 and a log2 fold-change cutoff of 2.5 were applied. C) PCA of mass-spec analysed pulldowns from HEK293T cells overexpressing empty vector (Control) or 2xHA-2xStrep-tagged AMPKα1. The plot shows PC1 and PC2 explaining 92.5% of the total variance. D) Venn diagram of overlapping proteins in OTULIN WT, OTULIN ΔPBM, and AMPK interactomes. The number of overlapping proteins and the percentage of total significantly upregulated proteins are indicated.

**Supplementary Figure S2. Relating to Figure 2.**
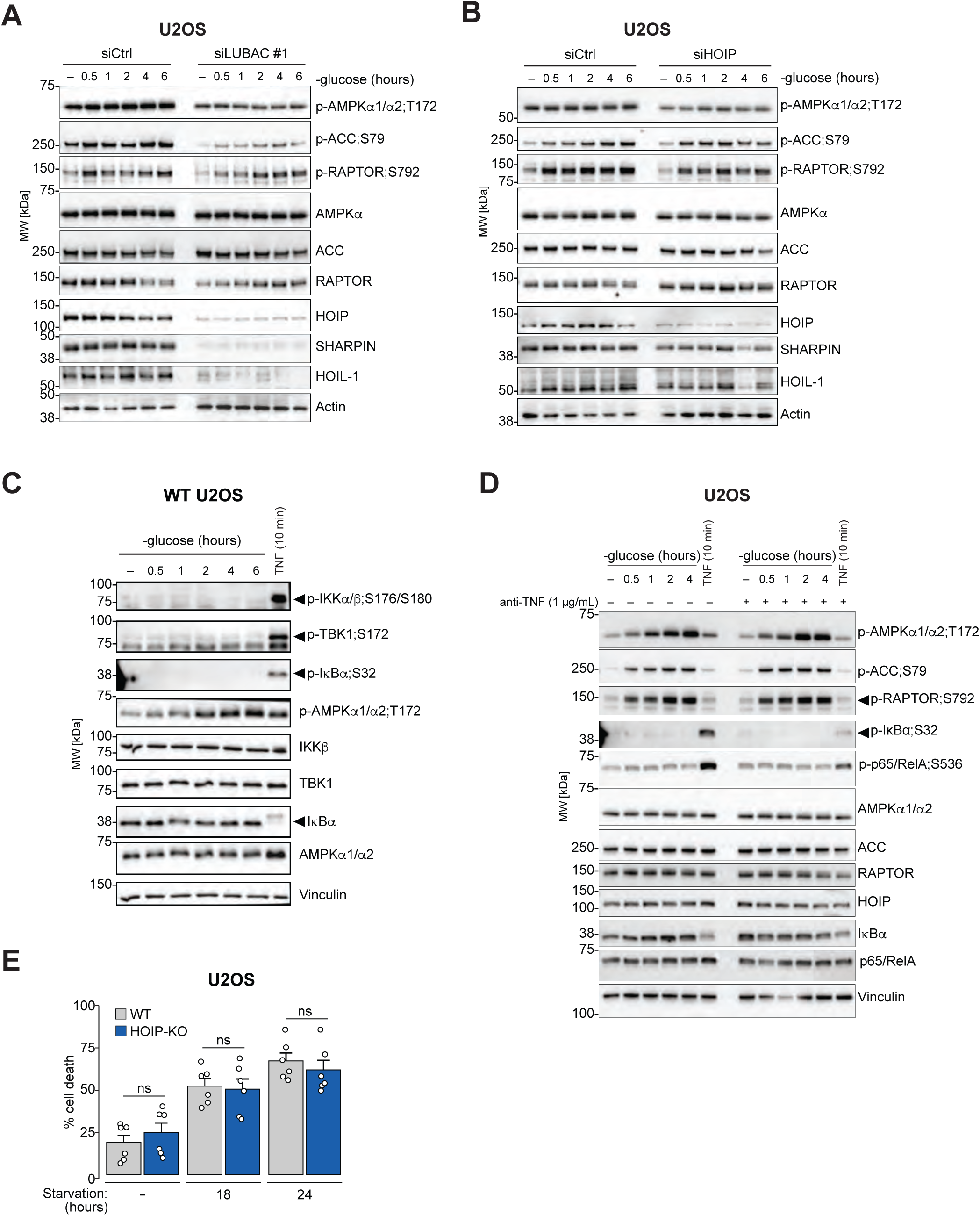
A) U2OS cells were transfected with siRNAs targeting HOIP (oligo #1), HOIL-1, and SHARPIN (LUBAC #1) or control (siCtrl) and starved in glucose-free medium as indicated. Samples were analysed by immunoblotting. B) U2OS cells transfected with siRNAs targeting HOIP alone (oligo #1) were treated and analysed as in (A). C) U2OS cells were starved in glucose-free medium as indicated or treated with 10 ng/mL TNF for 10 minutes and analysed by immunoblotting. D) U2OS cells were treated as in (A) with pre-treatment for 60 minutes with a TNF-blocking antibody (anti-TNF), and samples were analysed by immunoblotting. E) Cell death in U2OS WT (grey) or HOIP-KO cells (blue) at 0, 18, and 24 hours of glucose starvation. Cell death is expressed as a percentage of the maximal signal achieved by lysing cells with lysis buffer. Data are combined from two biologically independent replicates, with three replicates per treatment. A two-sided Student’s *t*-test was used to determine statistical significance. ns, non-significant. All blots show a single experiment representative of three biologically independent replicates.

**Supplementary Figure S3. Relating to Figure 3.**
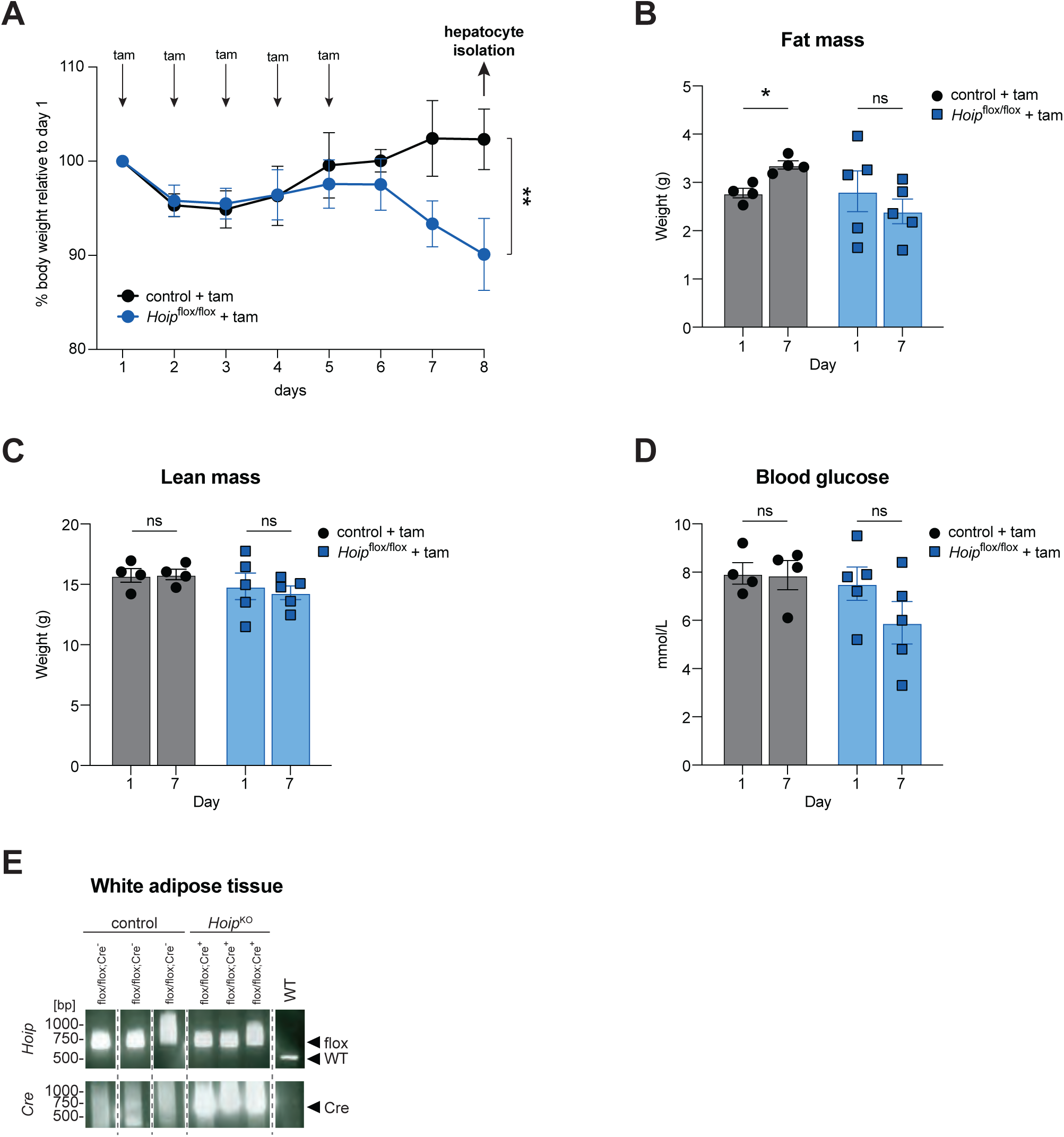
A) Body weight of control (black) and *Hoip*^flox/flox^ (blue) 10–14-week-old female mice treated with tamoxifen (tam) for the five days, shown as a percentage of body weight relative to day one. Hepatocyte isolation was carried out on day eight. Dots show the mean of 6 (control) or 7 (*Hoip*^flox/flox^) individual mice ± SEM. Statistical testing was performed by unpaired Students *t*-test on day 8 data points. ns, not significant; **p<0.01. B) Fat mass measured in grams by MRI in control and *Hoip*^flox/flox^ mice on day one and day seven after tamoxifen (tam) treatment. Each dot represents one mouse (n=4 or n=5 per group). Statistical testing was performed by paired Students *t*-test. ns, not significant; *p<0.05. C) Same as (B), but for lean mass. D) Blood glucose expressed as mmol/L in control and *Hoip*^flox/flox^ mice on day one and day seven of the experiment. Statistical testing was performed by paired student’s *t*-test. ns, not significant. E) Genotyping of mice used for the isolation of subcutaneous white adipose tissue for the presence of floxed *Hoip* alleles and the Cre recombinase (Cre) transgene. PCR reactions show the presence of floxed *Hoip* alleles in all animals (lanes 1–6), indicated by a band at the expected size (650 bp). Lanes 4–6 correspond to knockout (KO) animals, which also show the Cre transgene band. Lane 7 shows a WT genotype with *Hoip* at 507 bp and no Cre transgene.

**Supplementary Figure S4. Relating to Figure 4.**
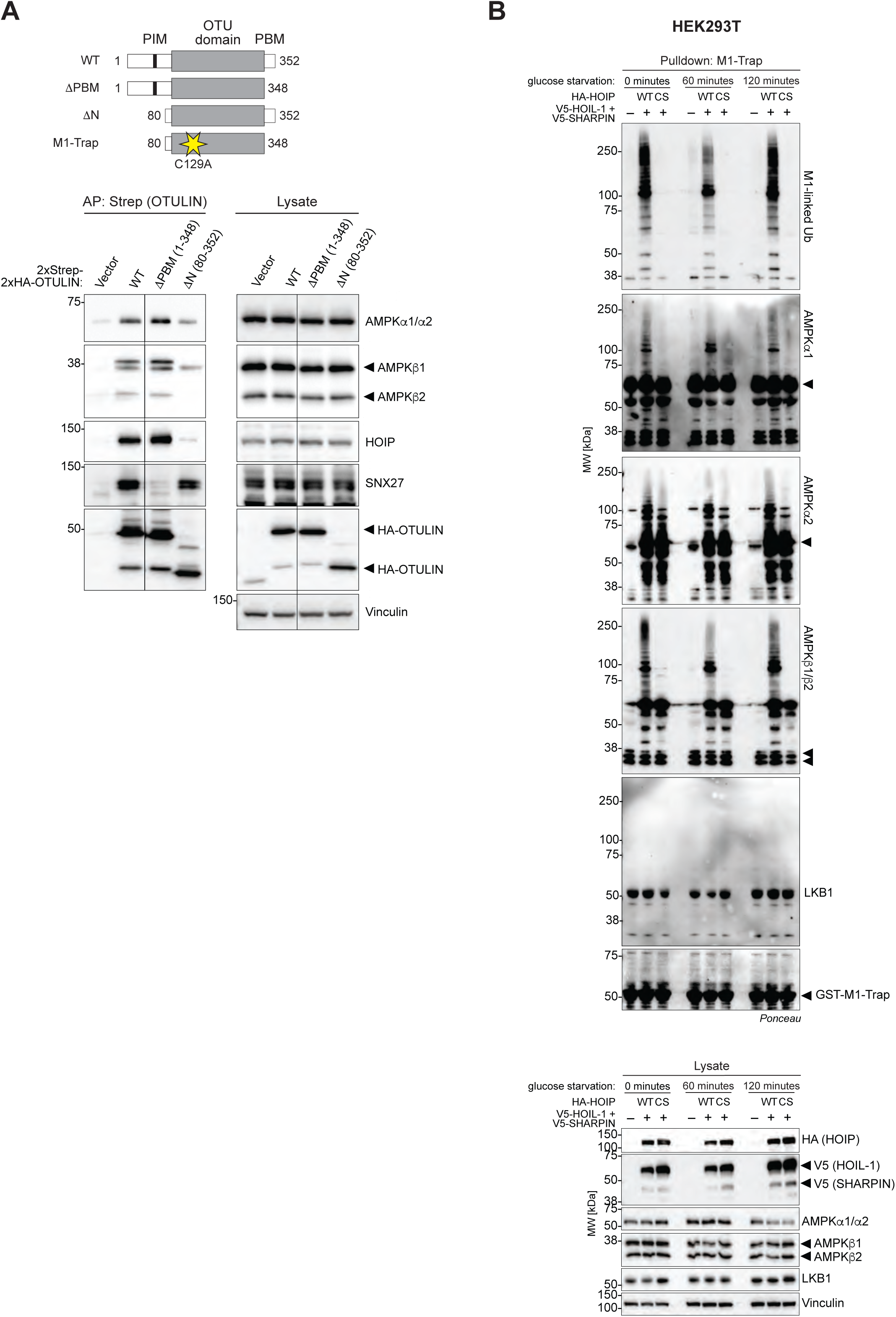
A) Top: Schematic representation of WT, PBM-deleted (ΔPBM), N-terminus deleted (ΔN) OTULIN, and the M1-Trap construct. Bottom: Strep-tagged OTULIN was affinity-precipitated from lysates of HEK293T cells transfected with empty vector or 2xHA-2xStrep-OTULIN variants as indicated. Samples were analysed for co-purified proteins by immunoblotting. Vertical lines indicate where blots from the same membrane have been cut and stitched. B) HEK293T cells were transfected with empty vector or LUBAC components HOIL-1, SHARPIN, and WT or catalytically inactive HOIP C885S and starved in glucose-free medium for 60 or 120 minutes. Endogenous M1-linked Ub chains were affinity-precipitated using recombinant GST-M1-Trap, and samples were analysed by immunoblotting. Blots from a single experiment representative of three biologically independent replicates are shown.

**Supplementary Figure S5. Relating to Figure 5.**
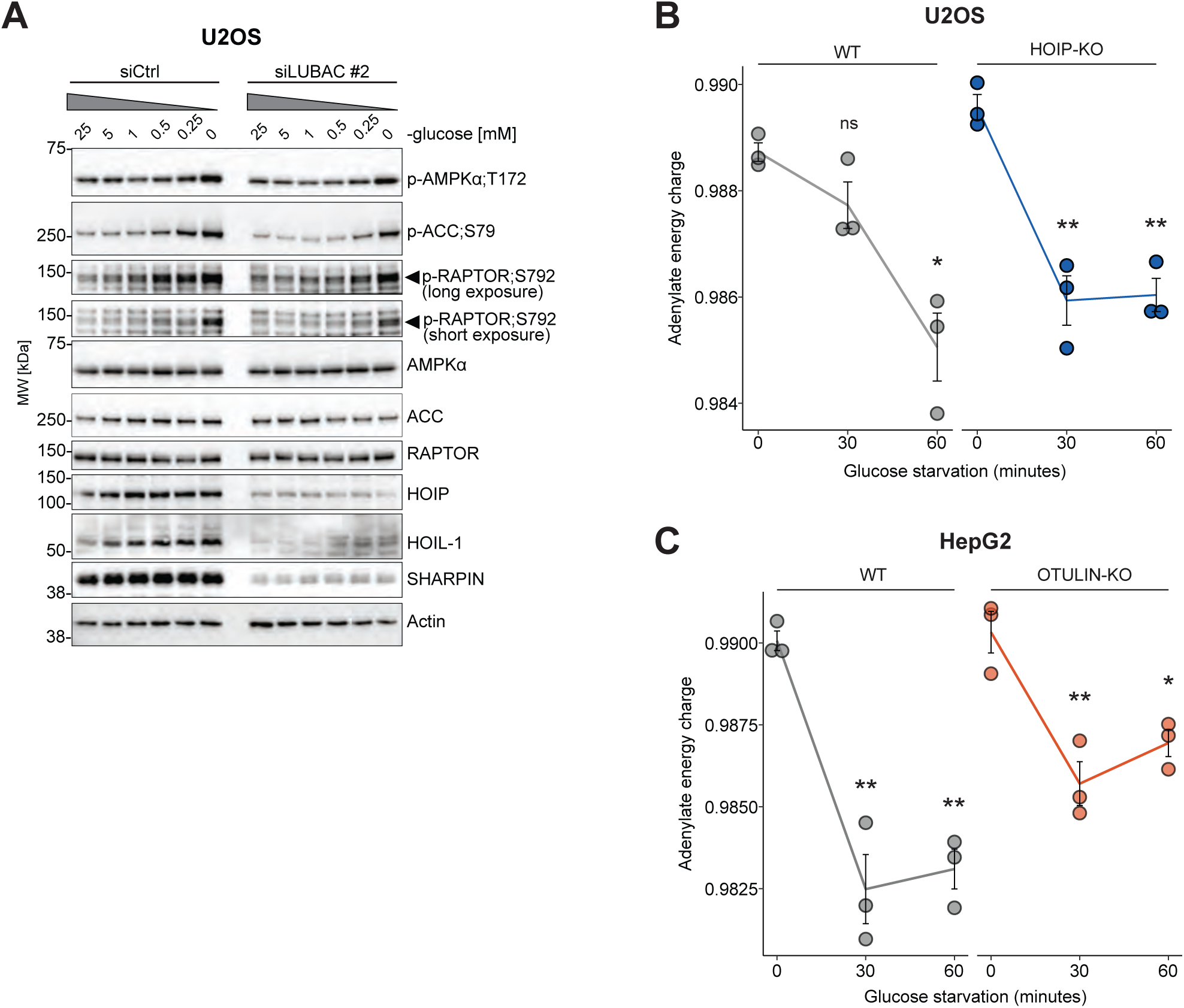
A) U2OS cells transfected with siRNAs targeting HOIP (oligo #2), HOIL-1, and SHARPIN (LUBAC #2) or control (siCtrl) were cultured in medium containing gradually lower concentrations of glucose for 60 minutes and analysed by immunoblotting. B-C) U2OS WT and HOIP-KO (B) or HepG2 WT and OTULIN-KO (C) cells were starved in glucose-free medium as indicated, followed by nucleotide extraction. AMP, ADP, and ATP levels were quantified using peak areas determined by LC-MS/MS and converted to adenylate energy charge (AEC). The plot shows mean AEC ± SEM for three independent experiments. Two-sided Student’s *t*-test was used to determine statistical significance. ns, not significant; *p<0.05; **p< 0.01.

**Supplementary Figure S6. Relating to Figure 6.**
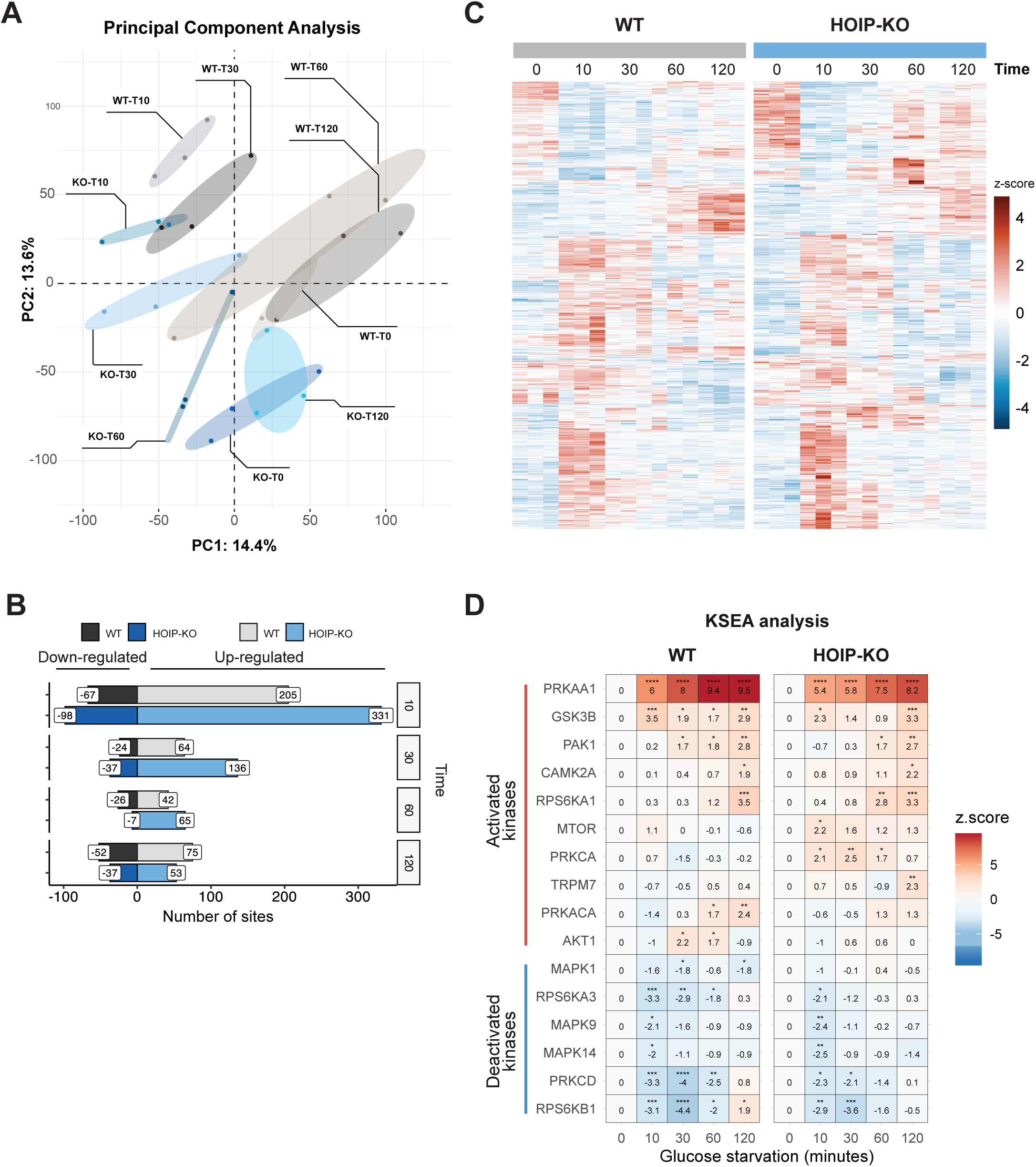
A) Principal Component (PC) Analysis of the samples used in the phosphoproteomic analysis. U2OS WT (grey hues) and HOIP-KO cells (blue hues) were deprived of glucose for 0, 10, 30, 60, or 120 minutes in triplicate. The plot shows the clustering of biological replicates in PC1 and PC2 dimensions, which explain 28% of the total variance. B) Bar plot showing the number of differentially altered phosphosites in WT (grey hues) and HOIP-KO cells (blue hues) at different time points after glucose starvation. Numbers of significantly upregulated sites are shown to the right side of 0 on the X-axis, and numbers of significantly downregulated sites are shown to the left side. Numbers are indicated at the top of the bars. Significant sites were determined by BH-adjusted FDR<0.05 and fold-change (FC) > 1.5. C) Heatmap of normalised phosphosite intensities for all sites shown in (B) clustered using hierarchical clustering based on correlation. D) Heatmap of kinase activity scores in WT and HOIP-KO cells over time, inferred using the Kinase-Substrate Enrichment Analysis (KSEA) tool. Normalised abundances of all phosphosites belonging to the “*Slow up*” cluster from Figure 6C were used to infer kinase activities. The colour intensity indicates the KSEA z-score which is also reported in numbers. Higher scores imply more kinase activity. The top 10 most active kinases in WT and HOIP-KO are shown. ns, not significant; *p<0.05; **p< 0.01; ***p<0.001; ****p<0.0001.

**Supplementary Figure S7. Relating to Figure 7.**
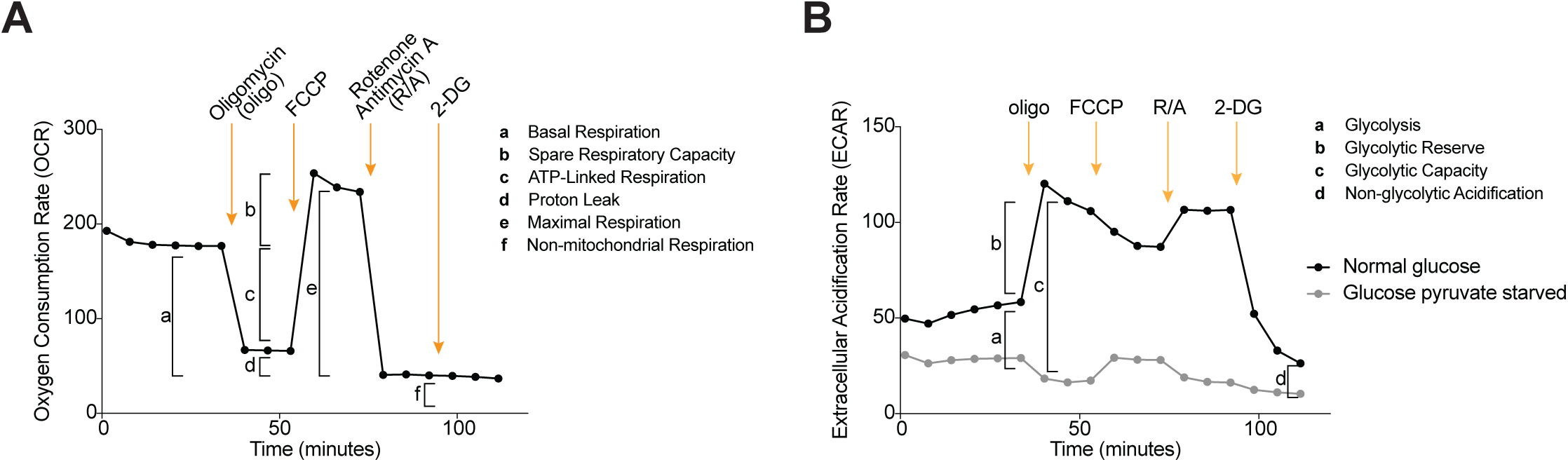
(A) Oxygen consumption rate (OCR) was quantified over time using the Seahorse extracellular flux (XF) assay. Basal respiration was calculated as the difference between the average OCR measured before injection of inhibitors, and the final OCR measurement after rotenone/antimycin A (R/A) injection, which represents only non-mitochondrial respiration. Maximal respiration was determined as the difference between the maximum OCR recorded after FCCP injection—which collapses the mitochondrial proton gradient, forcing complex IV consumption to reach its maximum—and non-mitochondrial respiration. Spare respiratory capacity was calculated as the difference between maximal respiration and the average OCR measured in the first six recordings. ATP-linked respiration was computed as the difference in average OCR before and after oligomycin injection, a direct inhibitor of complex V and ATP-linked respiration. Proton leak was calculated as the difference between the average OCR measured after oligomycin injection and non-mitochondrial respiration. (B) Extracellular acidification rate (ECAR) was quantified over time using the Seahorse extracellular flux (XF) assay. Basal glycolysis was calculated as the difference between the average ECAR measured before injection of inhibitors, and the last ECAR measurement after 2-deoxyglucose (2-DG) injection, which with directly glycolysis inhibited, represents non-glycolytic acidification. Glycolytic capacity was determined as the difference between the maximum ECAR after oligomycin injection, which forces cells to shift to glycolysis by inhibiting ATP synthase, and non-glycolytic acidification. Glycolytic reserve was calculated as the difference between glycolytic capacity and the average ECAR measured in the first six recordings. Prior to the assay, cells were incubated for 1 hour in media devoid of glucose and sodium pyruvate (referred to as glucose starvation or starved cells) to stimulate AMPK ac

## Methods

### Cell lines and primary cultures

AML12, HEK293T, HepG2, and U2OS, and mouse embryonic fibroblast (MEF) cell lines, as well as primary hepatocytes, were cultured in Dulbecco’s Modified Eagle’s Medium (DMEM) with high glucose, GlutaMAX™, and pyruvate (Invitrogen, 31966047) supplemented with 10% (v/v) foetal bovine serum (FBS) (Biowest, S181B) and 1% (v/v) penicillin-streptomycin 10,000 U/mL (Gibco, L0022). U2OS cells with double knockout (DKO) of AMPKa1 and AMPKa2^87^ as well as WT MEFs and MEFs with knock-in of HOIP-C879S^5^ have been described previously. All cell lines were kept in mycoplasma-free culture and tested regularly for contamination. Human fibroblasts derived from an ORAS patient (female) harbouring the OTULIN^G281R^ mutation and a healthy unrelated donor (male) (previously described^16^) were cultured in Ham’s F-12 Nutrient Mix supplemented with 20% FBS and 1% PS. Written informed consent was obtained from all subjects and family members. The study was approved by the ethical committee of Hadassah Medical Center and the Ministry of Health, Israel. All cells were cultured at 37°C in humidified incubators with 5% CO_2_ pressure.

### Mouse strains for isolation of primary hepatocytes in *Hoip*^flox/flox^ mice

The C57BL/6N-Rnf31^<tm1c(KOMP)Wtsi>/Tcp^ mouse line was made as part of the NorCOMM2 project with C57BL/6N-Rnf31^<tm1a(KOMP)Wtsi>/Tcp^ made from KOMP ES cells^88^ at the Toronto Centre for Phenogenomics. It was obtained from the Canadian Mouse Mutant Repository and was rederived on C57BL/6N background at the SPF facility at the University of Copenhagen, Faculty for Health and Medical Sciences. The B6.129-*Gt(ROSA)26Sor*^tm1(cre/ERT2)Tyj/J^ were from the University of Copenhagen transgenic mouse library and were similarly rederived on C57BL/6N background at the same SPF facility. All mouse strains were bred under SPF conditions and all breeding and mouse experiments were approved by the Danish Animal Experiments Inspectorate. Tamoxifen was dissolved in vehicle of 10% absolute ethanol and 90% sunflower seed oil at 20 mg /mL and was administered to mice by intraperitoneal injection (75 mg/kg bodyweight) every day for 5 days to induce deletion of *Rnf31*; control mice were injected with equal volume of vehicle only. Body weight was monitored daily for seven days. Whole-body composition analysis was conducted by TD-NMR (Minispec LF90II, Bruker; Billerica, MA, USA). *Ad lib*-fed blood glucose was measured once before anaesthesia for hepatocyte isolation using a handheld glucometer (Contour NEXT, Bayer), and euthanasia was confirmed by cervical dislocation. Agarose gel electrophoresis was used to genotype mouse ear DNA for the presence of WT and mutant alleles. Genomic DNA was extracted and amplified using allele-specific primers. PCR products were separated on a 1% agarose gel stained with SYBR Safe, and bands were visualized under UV light. *Hoip*^flox/flox^-CreERT2^+^ mice were used in experiments with *Hoip*^flox/flox^- or *Hoip*^WT/flox^-CreERT2^-^ animals as controls.

### Mouse strains for analysis of white adipose tissue in *Hoip*^A-KO^ mice

Mice with adipocyte-specific deletion of *Hoip (Hoip*^A-KO^) were generated by crossing the *Hoip*^flox/flox^ mice^89^ with an AdipoQ.cre Strain #:010803 from The Jackson Laboratories^90^. Cre^-^ littermates were used as control. 6-7-week-old male mice were bred and fed with a normal diet (DIO LS, 13% fat, 11% sucrose; Ssniff, E15748-04) for 16 weeks simultaneously using the same parental line. 3-5 mice were placed per cage and body weight gain was measured weekly. At the end of the feeding protocol, mice were euthanized using Ketamine/Xylazine Cocktail (100 mg/kg Ketamine and 20 mg/kg Xylazine) with a dose of 5 µL/g, followed by heart puncture, cervical dislocation and tissue harvesting. All animal experiments were approved by the German Federal Ministry for Nature, Environment and Consumers’ Protection of the state of North Rhine-Westphalia and were performed by the respective national, federal and institutional regulations. Agarose gel electrophoresis was used to genotype mouse ear DNA for the presence of WT and mutant alleles. Genomic DNA was extracted and amplified using allele-specific primers. PCR products were separated on a 2% agarose gel stained with SYBR Safe, and bands were visualized under UV light. *Hoip*^flox/flox^- *Adipoq*Cre^+^ mice were used in experiments with *Hoip*^flox/flox^-*Adipoq*Cre^-^ animals as controls.

### Drosophila melanogaster husbandry

*Drosophila melanogaster* flies were maintained at 25 °C with a 12-h light–dark cycle on medium containing agar 0.6% (w/v), malt 6.5% (w/v), semolina 3.2% (w/v), baker’s yeast 1.8% (w/v), nipagin 2.4%, and propionic acid 0.7%. Fly stocks used in this study: Daughterless-Gal4 (referred to as *DaGal4*) as control flies, Mi{ET1}LUBELMB00197 (referred to as *lubel^ΔRBR^*, Bloomington, #22725), UAS-LUBEL-RNAi P{GD7269} (referred to as *lubel^RNAi^*, Vienna Drosophila resource centre, #18055), UAS-CYLD-RNAi P{GD4739} (referred to as *cyld^RNAi^*, Vienna Drosophila resource centre, #15340), *cyld^MR4^* ^91^. UAS RNAi was driven by Daughterless-Gal4.

## METHOD DETAILS

### Plasmids and cloning

Desired inserts were PCR-amplified from in-house constructs or via reverse transcription (QuantiTect Reverse Transcription Kit, Qiagen) of mRNA isolated from HEK293T cells using the SV Total RNA Isolation System (Promega, Z3101). Purified PCR products were cloned into pcDNA3.1 or pSpCas9 vectors using In-Fusion primers and cloning kit (Takara Bio, 638948). Transformation was performed by heat-shocking Stellar™ Competent *E. coli* for 45 seconds at 40°C, followed by plating on LB agar containing ampicillin (100 µg/mL). Single colonies were picked and expanded in LB media with ampicillin (100 µg/mL), and plasmids were purified using the E.Z.N.A. Mini Kit I (Omega Bio-tek, D6942). Inserts were sequenced by Sanger sequencing (Macrogen Europe) using primers targeting the CMV immediate early promoter and a custom pcDNA3.1 reverse primer (sequences in KRT). Validated plasmid DNA was scaled up and purified for cell culture using the NucleoBond Xtra Midi kit (Macherey-Nagel, 740410).

### Plasmid DNA transfection

Cells were cultured until they reached approximately 80% confluency, at which point they were transfected with plasmid DNA to achieve transient overexpression using the X-tremeGENE HP DNA Transfection Reagent (Roche) following the manufacturer’s instructions. The transfection reagent-to-DNA ratio used was 2:1 (v/w). Cells were incubated for 24-48 hours before experiment onset.

### Generation of CRISPR knockout cells

For the generation of U2OS cells with CRISPR–Cas9 knockout of HOIP or OTULIN, and AML12 cells with knockout of *Hoip*, cells were grown to 60% confluency and then transfected with gRNA cloned into the pSpCas9(BB)-2A-Puro (PX459) backbone (Addgene, #62988) (sequences in KRT). 48 hours post-transfection, puromycin selection was initiated at a concentration of 5 µg/mL, with media being replaced daily for five days. Following selection, cells were plated into 96-well plates and grown from single clones. Successful knockout was confirmed via immunoblot analysis. Once confirmed, individual clones were maintained in selection-free media. OTULIN-deficient HepG2 cells were similarly generated using CRISPR–Cas9 gene targeting, with a sgRNA designed using the CRISPRscan tool (sequence in KRT, OTULIN gRNA #2) and cloned into the pSpCas9(BB)-2A-GFP (PX458) vector (Addgene, #48138). 1×10^6^ cells were electroporated with 10 μg DNA using the NEPA21 cuvette electroporator. Two days post-electroporation, GFP-positive single cells were sorted into 96-well plates. Clones were expanded and screened by immunoblotting to select OTULIN-knockout clones.

### siRNA transfection

Cells were seeded and cultured to 50-60% confluency before being transfected using Lipofectamine® RNAiMAX Transfection Reagent (Thermo Fisher Scientific) following the manufacturer’s instructions. The final concentration of siRNA was either 40 nm (for siOTULIN and siHOIP) or 50 nM (siLUBAC: 25 nM siHOIP + 12.5 nM siRBCK1 + 12.5 nM siSHARPIN). Non-targeting siRNA (siCtrl) was used as control at the same concentrations. Oligo sequences are listed in the KRT. After overnight incubation, the transfected cells were reseeded and allowed to grow for an additional 24 hours to facilitate target protein depletion. Following this, the cells were used directly in the relevant assays.

### Cell treatments

For glucose starvation experiments, cells were washed with PBS and incubated in glucose-free DMEM (without sodium pyruvate), supplemented with 10% FBS and 1% Penicillin-Streptomycin. Control cells were similarly washed and incubated in fresh growth medium containing 4.5 g/L D-glucose. For Bafilomycin A1 treatment, cells were incubated either in regular growth medium, glucose-free DMEM, or Hank’s Balanced Salt Solution (HBSS) for 2 hours with or without 100 nM Bafilomycin A1 to block fusion of the autophagosome with lysosomes. For the protein stability assay, cells were treated with either 100 µM cycloheximide or DMSO for up to 6 hours, under both high glucose and glucose starvation conditions. NF-κB signalling was assessed by treating cells with 10 ng/mL human TNF-α for 10 minutes as a positive control. To block TNF signalling, cells were pre-treated with 1 µg/mL anti-TNF antibody (BioXCell, Adalimumab biosimilar, SIM0001) for 1 hour, followed by glucose starvation or 10 ng/mL TNF-α treatment with continuous TNFR blocking. All experiments were performed in triplicate.

### Starvation and survival in *Drosophila* flies and larvae

To assess survival during starvation, 20-30 adult flies per genotype were starved in tubes containing a layer of 2 % agarose for up to 48 hours and the survival was counted at indicated time points. Larvae were starved on 2% agarose for 3 hours before lysis and immunoblot analysis.

### Cell lysis and protein extraction for immunoblotting

Cells were scraped in lysis buffer supplemented with cOmplete™, EDTA-free Protease Inhibitor Cocktail (Roche, 11873580001), PhosSTOP™ (Roche, 4906845001), DUB inhibitor: 50 mM N-Ethylmaleimide (NEM), and 1 mM dithiothreitol (DTT). Lysates were cleared by centrifugation for 10 minutes at 13,000 rcf at 4°C. Protein concentration was measured with Bradford assay and concentrations were adjusted accordingly to ensure equal loading. Lysates were mixed with 4x Laemmli sample buffer (LSB) and heated for 5 min at 60°C.

### Tissue lysis and protein extraction for immunoblotting

Inguinal subcutaneous white fat pads were harvested from mice and snap-frozen in liquid nitrogen. Frozen tissue samples (approximately 100 mg) were cut on dry ice and placed in 2 mL round-bottom Eppendorf tubes containing 5 mm stainless steel beads. In a first step lysis buffer (50 mM Tris (pH 7.5), 150 mM NaCl, without detergent) supplemented with protease inhibitor, phosphatase inhibitor, NEM, DTT, and was added at a ratio of 5 μL/mg ensuring a minimum volume of 500 μL per sample. Tissues were homogenized using the TissueLyser II (Qiagen) for 3 min at 25 Hz. The homogenization process was repeated until tissues were fully disrupted and homogeneous. Samples were centrifuged at 6,000 rcf for 15 min at 4°C to separate the lipid layer. The fat cake (white lipid layer) was carefully removed, and the NP-40 was added to the remaining solution at 0.5% v/v and incubated for 30 minutes on ice. A final centrifugation at 13,000 rcf for 15 min at 4°C was performed to remove residual lipids. The clarified lysates were transferred to new 1.5 mL tubes and stored at -80°C until further use.

Whole liver lysates from 9-day-old control and *Otulin*^Δhep^ mice were prepared as previously described^28^ using a TissueLyser II (Qiagen) to lyse tissue samples in RIPA buffer (50 mM Tris pH 7.4, 1% NP-40, 0.5% deoxycholate, 0.1% SDS, 150 mM NaCl, 2 mM EDTA, and 5 mM MgCl2) supplemented with complete protease inhibitor cocktail and PhosSTOP phosphatase inhibitors (Roche).

Larvae from *Drosophila melanogaster*: Analysis of phospho-AMPK levels in *Drosophila melanogaster* was done in third instar larvae (10 larvae/genotype). The larvae were homogenized and lysed 15 min on ice in lysis buffer (50mM Tris (pH 7.5), 150mM NaCl, 1% Triton X-100, 1mM EDTA (pH 8) and 10% glycerol) supplemented with Pierce™ Protease Inhibitor (Thermo Fisher Scientific). The lysates were cleared before the addition of LSB and heated for 5 min.

### Immunoprecipitation and affinity-precipitation

HEK293T or U2OS cells were transfected and treated as indicated. Cells were washed once with ice-cold PBS and lysed on ice using IP buffer (25 mM HEPES (pH 7.4), 150 mM KCl, 2 mM MgCl₂, 1 mM EGTA, 0.5% Triton X-100) supplemented with protease, phosphatase, and DUB inhibitors. Lysates were cleared by centrifugation at 13,000 rpm for 10 minutes at 4°C, and supernatants were transferred to clean tubes and supplemented with 1 mM DTT. For endogenous IPs, primary antibodies (concentrations according to manufacturers’ recommendations) were added along with 20 µl bead slur/sample pre-washed protein A/G agarose beads (Pierce, 20423) or magnetic beads (Pierce, 88803) and incubated with lysates at 4°C for 2–18 hours with gentle agitation. For tagged protein immunoprecipitation, lysates were incubated with either 20 µl bead slur/sample pre-washed magnetic anti-HA beads (Pierce, 13474229) or pre-washed magnetic anti-FLAG^®^ M2 beads (Millipore, M8823). For affinity-precipitation lysates were incubated with 40 µl bead slur/sample of pre-washed Strep-Tactin™XT Superflow™ (IBA Lifesciences, 15583756). Beads were washed two times with 500 µL of ice-cold high-salt IP buffer (IP buffer with 1 M KCl) followed by two times with 500 µL of ice-cold regular IP buffer. After incubation and washing, bound material was eluted in either LSB (for SDS-PAGE and immunoblotting) by heating to 60°C for 5 minutes, or in MS lysis buffer (6 M guanidinium hydrochloride, 10 mM TCEP, 40 mM CAA, 50 mM HEPES, pH 8.5) for mass spectrometry analysis.

### Purification of Endogenous Ubiquitin Conjugates

Endogenous Ub conjugates were purified using either the M1-SUB or the M1-Trap (see KRT). Cells (from a 15 cm dish of U2OS or HEK293T, depending on the experiment) were lysed in 500-1000 µL of ice-cold TUBE buffer (20 mM Na₂HPO₄, 20 mM NaH₂PO₄, 1% NP-40, 2 mM EDTA) supplemented with protease inhibitor, phosphatase inhibitor and NEM. Lysates were cleared at 13,000 rcf for 10 minutes at 4°C, and the supernatants were transferred to clean tubes. Samples were supplemented with 1 mM DTT and 100-150 µg/mL of purified GST-M1-SUB or GST-M1-Trap and incubated with 25-40 µL bead slurry/sample of pre-washed Glutathione Sepharose 4B (Sigma, GE17-0756-01) or Glutathione Magnetic Agarose Beads (Millipore, G0924). Samples were incubated for 2-18 hours at 4°C with gentle agitation, washed three times with ice-cold PBS-T, and eluted using LSB for immunoblot analysis.

### SDS-PAGE and immunoblotting

For SDS-PAGE and immunoblotting, samples were resolved on NuPAGE Tris-Acetate 3–8% or Bis-Tris 4–12% gels according to manufacturer instructions, or self-cast 9% Bis-Tris gels. Proteins were transferred to 0.2 µm nitrocellulose membranes using the Trans-Blot Turbo system. Membranes were stained with Ponceau S to verify transfer efficiency and equal loading, then blocked in 5% milk in PBS-T (0.1% Tween-20 in PBS). After washing with PBS-T, membranes were incubated overnight at 4°C with primary antibodies (1:1000 dilution in 3% BSA, 0.01% NaN₃ in PBS-T). The following day, membranes were washed three times with PBS-T and incubated with HRP-conjugated secondary antibodies (see KRT) for 45 minutes at room temperature. After additional washes, blots were developed using Clarity™ Western ECL substrate (Bio-Rad, 1705061) and imaged on a Bio-Rad ChemiDoc™ Imager. Image Lab software (Bio-Rad) was used for image assembly and analysis.

### Immunoprecipitation-coupled kinase assays

For immunoprecipitation-coupled kinase assays, cells were lysed in lysis buffer (50 mM Tris–HCl (pH 7.4), 150 mM NaCl, 1% (v/v) Triton X-100, 50 mM NaF, 5 mM sodium pyrophosphate, 1 mM EDTA, 1 mM EGTA, 1 mM DTT, 0.1 mM benzamidine, 0.1 mM PMSF). Lysates were cleared and protein concentration was determined using Bradford assay. Equal amount of anti-AMPK-α1 and -α2 antibodies were non-covalently coupled to protein G-Sepharose beads. The beads were incubated with 100-300 µg of cell lysates on a roller mixer for 2 hours at 2–8°C to immunoprecipitate AMPK. Beads were washed twice with high salt IP buffer (IP buffer with 1 M NaCl), twice with regular IP buffer, and twice with HEPES assay buffer (50 mM HEPES, 0.02% Brij-35, and 1 mM DTT added fresh). After washing, beads were resuspended in HEPES assay buffer and aliquoted into 1.5 mL Eppendorf tubes to a final volume of 20 µL for subsequent kinase assays. AMPK activity was measured based on its ability to phosphorylate the synthetic AMARA peptide (AMARAASAAALARRR), derived from rat acetyl-CoA carboxylase (ACC), a known AMPK substrate^92^. The kinase reaction was initiated by adding 30 µL of HEPES assay buffer containing 200 µM AMP, 200 µM [γ-^33^P]-ATP (Hartmann Analytic), 5 mM MgCl₂, and 200 µM AMARA peptide to a final volume of 50 µL. Assays were incubated at 30°C on an orbital shaker for 15 minutes. The reaction was stopped by spotting 30 µL of the mixture onto P81 filter paper, followed by immersion in 1% orthophosphoric acid. Filter papers were washed with water to remove unincorporated ATP, dried at room temperature, and submersed in scintillation fluid. Radioactivity was measured on a scintillation counter. One unit of AMPK activity was defined as the incorporation of 1 nanomole of ^33^P into the AMARA peptide mg^-1^ min^-1^.

### Bioenergetics by Seahorse XF measurements

U2OS were seeded into XF96 Polystyrene Cell Culture Plate (Agilent Technologies) 24 to 48 hours before assay, at 30,000 cells per well. On the day of the assay, the assay medium was prepared fresh using Seahorse XF base medium (Agilent Technologies) adjusted to pH 7.4, supplemented with 2mM L-glutamine (Thermo Fisher Scientific) and optionally supplemented with 10mM glucose (Merck) and 1mM sodium pyruvate (Merck). At 1 hour before the assay, cells were incubated with assay medium, either present or absent for glucose and sodium pyruvate, then incubated at 37°C in an incubator absent of CO_2_. Six baseline OCR and ECAR recordings were made, followed by sequential injection of ATP synthase inhibitor oligomycin (Oligo; Merck 1μM), mitochondrial uncoupler, carbonyl cyanide p-triflouromethoxyphenylhydrazone (FCCP; Merck, 1μM), the Complex I and III inhibitors rotenone and antimycin A (R/A; Merck, 5μM) and the glucose analogue 2-deoxyglucose (2-DG; Merck, 50mM). Mitochondrial respiration and glycolytic activity of U2OS were measured using the Seahorse XFe96 Analyzer (Agilent Technologies) with downstream analysis using the Seahorse Wave Desktop Software (Agilent Technologies). Measurements were normalised to total well protein content using the Pierce™ BCA Protein Assay Kit (Thermo Fisher Scientific).

### Cell death assay

Cell death due to starvation was measured using the CellTox™ Green Cytotoxicity Assay (Promega, #G8741). Cells were seeded in 96-well plates 24 hours before the longest starvation timepoint, with seeding density adjusted based on the assumption of no proliferation during starvation and a doubling time of 20–24 hours. At the experimental endpoint, wells were washed once with PBS, followed by a 30-minute incubation in 100 µL of Assay Buffer containing CellTox™ Green Dye (1:1000), protected from light. Lysis buffer was added to positive control wells to ensure complete dye permeability. This control served as the maximum signal achievable from both proliferating and non-proliferating cells at the end of the experiment. Fluorescence was measured at 485–500nm Ex/520–530nm Em using a Victor Nivo plate reader (Perkin Elmer). Cell death was expressed as a percentage of the average fluorescent signal relative to the positive control.

### Purification of Ubiquitin Affinity reagents

M1-SUB (GST-NEMO-CoZi-His), and M1-Trap (GST-OTU-His, OTULIN aa 80-348, C129A) were purified from *E. Coli* BL21(DE3). Chemically transformed bacteria in LB-broth were grown to OD 0.6, induced with 0.5 mM IPTG and expressed overnight at 18°C. Bacteria were harvested, pelleted and flash frozen. Recombinant proteins were purified by IMAC or GST purification. For IMAC, pellets were resuspended in Buffer A (25 mM Tris, 200 mM NaCl, 20 mM Imidazole (pH 8.0)) supplemented with protease inhibitor cocktail (Sigma) and 10 U/ml rDNase I (Roche). After sonication, pelleting (29,000 rcf, 20 min, 4°C) and filtration (1 μm PES), the cleared lysate was loaded on a HisTrap FF Column (Cytiva). Protein was eluted by gradient up to 100% Buffer B (25 mM Tris, 200 mM NaCl, 500 mM Imidazole (pH 8.0)). Protein was further purified and buffer-exchanged into PBS (10 mM P_i_, 138 mM NaCl, 2.7 mM KCl (pH 7.4)) on a Superdex 75 Increase 10/300 GL SEC column (Cytiva) for further use. For GST purification, pellets were resuspended in PBS (see above), supplemented with protease inhibitor cocktail (Sigma) and 10 U/ml rDNase I (Roche). After obtaining the cleared lysate as described above, it was incubated with 0.5 mL slurry of Glutathione Sepharose 4B beads (Cytiva) at 4°C for 1 h while tumbling slowly in a 49 mL Econo-Column (BioRad). After releasing the flowthrough, beads were washed with ice cold PBS and eluted via addition of 5x 1 mL GSH-elution buffer (PBS supplemented with 50 mM GSH and 1 mM DTT). Purified protein was buffer-exchanged into PBS via NAP-25 columns (Cytiva) for further use.

### Isolation of primary hepatocytes

Murine hepatocytes were isolated using the method described by Gandin *et al*.^93^ with modifications. Briefly, the liver was perfused with washing buffer (10 mM HEPES and 200 µM EGTA in Hanks Buffered Saline Solution (HBSS without calcium or magnesium (Gibco, 14170112)) through the vena porta at a flow rate of 5 mL/min. The vena cava was cut to allow for drainage. This was followed by perfusion of digestion buffer containing collagenase from *Clostridium histolyticum* (Sigma, C5138) in HBSS with calcium and magnesium (Gibco, 24020117). The liver was then excised and transferred to a dish containing ice-cold plating medium (Medium-199 (Gibco, 31150-022) + 10% FBS, 1% (v/v) penicillin-streptomycin), and the gallbladder was removed. The resulting cell suspension was filtered through a 100 μm filter, centrifuged, and cells washed in plating medium. Viable hepatocytes were purified by centrifugation in 90% Percoll (Sigma P4937) in PBS. Live cells were washed in plating medium once more before being passed through a 70 μm filter. Finally, the cells were diluted in plating medium supplemented with 0.1 μm dexamethasone (Sigma, D4902) and 1 nM insulin (Sigma, I9278). Hepatocytes were seeded onto collagen-coated plates at a density of 200,000 cells/mL and allowed to attach for 2–4 hours before changing to culture medium (1g/L glucose DMEM Gibco 10567014). The following day, the hepatocytes were ready for experimental use. Glucose-free DMEM was used for hepatocyte starvation (Gibco, 11966025).

### Nucleotide extraction and measurement of ATP, ADP and AMP

Culture medium was aspirated from cells, followed by washing in ice-cold PBS and lysis in ice-cold 5% perchloric acid. Samples were vortexed and lysates clarified by centrifugation. The supernatant was collected and an equal volume of 1:1 mixture of tri-n-octylamine and 1,1,2-trichlorotrifluoroethane added. Samples were vortexed, centrifuged and the top (aqueous) phase collected. This procedure was repeated twice more, and nucleotides stored at -20°C before analysis. All centrifugation steps were performed at 13,000 rcf, 4°C for 3 minutes. The levels of AMP, ADP and ATP were measured using a TSQ Quantiva interfaced with Ultimate 3000 Liquid Chromatography system (Thermo-Scientific), as described by Ross *et al*. ^94^ will little modification of the LC gradient. The porous graphitic carbon column (Hyper Carb 30×1mm ID 3 mm; Part No: C-35003-031030, Thermo-Scientific) was maintained at a constant temperature of 42°C and equilibrated for 11 min with 7% buffer B at a constant flow rate 0.045 ml/min. Aliquot of 1 µl of each sample were loaded onto the column and the three nucleotides were eluted with a linear gradient of 7% to 12% buffer B over 3 min, then from 12%-90% buffer B within 2 min and finally from 90%-100 buffer B within 4 min. The Quantiva parameters and LC buffers were maintained as described by Ross *et al*. ^94^. The area of each nucleotide was determined by manually integrating area under the peak. The relative abundances of AMP, ADP and ATP were converted to adenylate energy charge (AEC) defined as:

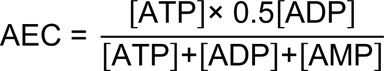

### Proteomics Sample preparation

Immunoprecipitated proteins were digested on-beads by eluting in 20 μL of MS lysis buffer (6 M Guanidinium-HCl, 10 mM TCEP, 40 mM CAA, 50 mM HEPES, pH 8.5). The samples were heated to 95°C for 5 minutes, followed by sonication at high intensity for 5 cycles of 30 seconds in a Bioruptor sonication water bath (Diagenode) at 4°C. Protein concentration was determined using the Pierce™ Rapid Gold BCA Protein Assay Kit (Thermo Fisher, A55860), and 500 μg of protein was used for digestion. Samples were diluted 1:3 with 10% acetonitrile in 50 mM HEPES (pH 8.5). Proteins were first digested with LysC (MS grade, Wako) at a 1:50 enzyme-to-protein ratio and incubated at 37°C for 4 hours. The samples were further diluted 1:10 with 10% acetonitrile in 50 mM HEPES (pH 8.5), and trypsin (MS grade, Promega) was added at a 1:100 enzyme-to-protein ratio for overnight digestion at 37°C. Digestion was quenched by adding 2% trifluoroacetic acid (TFA) to a final concentration of 1%. The resulting peptides were desalted using a SOLAμ™ SPE Plate (Thermo Fisher) according to the manufacturer’s instructions. Briefly, the sorbent was activated with 40 μL of 100% methanol (HPLC grade, Sigma) followed by 40 μL of Buffer B (80% acetonitrile, 0.1% formic acid). The plate was equilibrated twice with 40 μL of 1% TFA in 3% acetonitrile, after which the samples were loaded. The plates were washed twice with 200 μL of Buffer A (0.1% formic acid in water), and bound peptides were eluted into clean 500 μL Eppendorf tubes using Buffer A’ (40% acetonitrile, 0.1% formic acid). The eluted peptides were then concentrated using an Eppendorf Speedvac and reconstituted Buffer A* (0.1% formic acid + iRT peptides (Biognosys, KI-3002-2)). A total of 500 ng of peptides were loaded onto the instruments for analysis.

### MS data acquisition

Peptides were loaded onto a 2 cm C18 trap column (ThermoFisher, 164946), connected in-line to a 15 cm C18 reverse-phase analytical column (Thermo EasySpray, ES904) using 100% Buffer A at 750 bar, using the Thermo EasyLC 1200 HPLC system, and the column oven operating at 35°C. Peptides were eluted over a 70 minute gradient ranging from 6 to 60% of Buffer B at 250 nL/min, and the Q-Exactive instrument (Thermo Fisher Scientific) was run in a DD-MS2 top10 method. Full MS spectra were collected at a resolution of 70,000, with an AGC target of 3×10^6^ or maximum injection time of 20 ms and a scan range of 300–1750 m/z. The MS2 spectra were obtained at a resolution of 17,500, with an AGC target value of 1×10^6^ or maximum injection time of 60 ms, a normalised collision energy of 25 and an intensity threshold of 1.7e4. Dynamic exclusion was set to 60 s, and ions with a charge state <2 or unknown were excluded. MS performance was verified for consistency by running complex cell lysate quality control standards, and chromatography was monitored to check for reproducibility.

### Data analysis

All raw LC-MS/MS data files were processed together using Proteome Discoverer version 2.4 (Thermo Fisher). In the processing step, Oxidation (M), Acetylation (K), and protein N-termini acetylation and met-loss were set as dynamic modifications. All results were filtered with 0.05 delta Cn for PSMs. SequestHT was used as database, matching spectra against the *E.coli* database from Uniprot together with recombinant sequence. Identified proteins and their quantified abundances were imported into RStudio software (Posit, version 2024.09.0+375) and filtered for missing values, retaining proteins detected in at least 75% of bait samples (with no filtering on empty vector control samples). Abundances were normalized to total sample loading by scaling each row (protein) to the median of column (sample) sums divided by the total sample sum. After log2 transformation, missing values were imputed for “clean” hits using normal downshift (width = 0.3, downshift = 1.8). “Clean” hits are defined as those not detected in any replicate of control pulldowns but detected in ≥75% of replicates in bait pulldown. Heatmap was generated using the pheatmap package in R to visualize normalised protein intensities. Differentially enriched proteins were tested using limma linear modelling^95^ applying a fold-change cutoff of 2.5 and a false discovery rate (FDR) below 0.05 after Benjamin-Hochberg (BH)-adjustment for multiple testing. Proteins belonging to the KEGG pathway “hsa04152_AMPK_signaling_pathway” were extracted using the ‘EnrichmentBrowser’ package (version 2.34.1)^96^.

### Phosphoproteomics

#### Sample preparation

Proteins were extracted by lysing cells in 6 M GdnHCl in 250 mM HEPES (pH 8.0) using bead beater, reduced with 15 mM TCEP for 1h at 55°C, and alkylated with 30 mM IAA for 30 min at room temperature. After acetone precipitation, proteins were resuspended in small volume of 2.5 M GdnHCl in 100 mM HEPES (pH 8.0) and 120 µg protein of each sample were digested overnight with LysC and trypsin (ratio 50:1, final volume of 100 µL, final GdnHCl concentration of 0.75 M).

#### Enrichment of phospho-peptides and TMT labelling for proteomics

TMTpro™ 16-plex labelling was performed according to the manufacturer’s recommendations (Thermo Fisher, A44520). Two TMT 16-plexes were used for the 30 samples thus resulting in two batches. After combining same amount of each TMT channel, samples were acidified using 50% TFA (final concentration app. 0.5%, (pH<2.0) and peptides were purified by SPE using HR-X columns in combination with C18 cartridges (Macherey-Nagel): Columns were washed in Buffer A and eluted in Buffer B. Elutes were frozen in liquid nitrogen and lyophilized overnight. Peptide mixture was fractionated by HpH reverse phase chromatography resulting in eight fractions^97^. Phosphopeptide enrichment was done on each fraction according to Post *et al*.^98^ using Fe(III) cartridges on Bravo liquid handling system (Agilent). Flow-through was collected for global proteome analysis.

#### MS data acquisition

LC-MS/MS measurements were performed on an Exploris 480 mass spectrometer (Thermo Scientific) coupled to an EasyLC 1200 nanoflow-HPLC (Thermo Scientific). Peptides were separated on a fused silica HPLC-column tip (I.D. 75 μm, New Objective, self-packed with ReproSil-Pur 120 C18-AQ, 1.9 μm (Dr. Maisch) to a length of 20 cm) using a gradient of Buffer A and Buffer B: samples were loaded with 0% Buffer B with a flow rate of 600 nL/min; peptides were separated by 7%–38% B within 122 min with a flow rate of 250 nL/min. Spray voltage was set to 2.3 kV and the ion-transfer tube temperature to 250°C; no sheath and auxiliary gas were used. Mass spectrometers were operated in the data-dependent mode; after each MS scan (mass range m/z = 370 – 1750; resolution: 120’000) a maximum of twenty MS/MS scans were performed using an isolation window of 0.7, a normalized collision energy of 32%, a target AGC of 50% and a resolution of 45’000. MS raw files were analysed using ProteomeDiscoverer (version 2.5, Thermo Scientific) using a Uniprot full-length Yeast database and common contaminants such as keratins and enzymes used for digestion as reference. Carbamidomethyl cysteine (and phospho-STY for phospho-enriched samples) was set as fixed modifications and protein amino-terminal acetylation, and oxidation of methionine were set as variable modifications. The MS/MS tolerance was set to 0.6 Da and three missed cleavages were allowed using trypsin/P as enzyme specificity.

#### Data analysis

Identified peptides and proteins, along with their quantified abundances from Proteome Discoverer, were imported into R software (Posit). The intensities of phosphosites on peptides were summed, accounting for different phosphorylation multiplicities. The datasets were filtered for missing values to retain only the peptides and proteins detected in all samples (across the two TMTplexes). Protein group abundances were used for the proteome analysis, while phosphopeptide abundances were used for the phosphoproteome. Data normalization was carried out in three steps for both proteome and phosphoproteome datasets. First, sample loading was adjusted by multiplying the abundances by a sample-specific scaling factor. Subsequently, artificial internal reference standards were generated to create protein/peptide-specific scaling factors for the two TMTplex batches. Finally, phosphosite abundances were corrected by multiplying them with their corresponding protein abundance scaling factor. In the case of phosphosites without a corresponding protein value, the scaling factor was set to 1. After log2 transformation, differentially regulated phosphosites were tested by limma linear modelling ^99^, applying a fold-change cutoff of 1.52 (log2-FC=0.6) and a BH-adjusted false discovery rate (FDR) below 0.05. Heatmap was generated using the pheatmap package in R to visualise normalised phosphosite intensities.

To identify patterns of phosphorylation dynamics, fuzzy c-means clustering was employed using the Mfuzz R package^100^. Before clustering, normalised phosphosite abundances were scaled. Clustering was conducted with optimal parameters n=4 and c=1.99, which were determined using the built-in functions of the package. Kinase-substrate enrichment analysis was conducted using the KSEA App R package ^75^ using annotations from both PhosphoSitePlus (July 2016 release) and NetworKIN with a minimum NetworKIN score of 10 and otherwise default parameters. For functional enrichment, relevant phosphosites were collapsed to their UNIPROT protein ID and sorted by fold-change in descending order. Gene-Set Enrichment Analysis (GSEA) was performed using the clusterProfiler R package ^101^. The human proteome served as the background for the representation analysis. KEGG pathway gene sets were tested for enrichment with a BH-adjusted p-value cutoff of <0.05 or 0.01 as indicated.

## QUANTIFICATION AND STATISTICAL ANALYSIS

### Statistical testing

Data are presented as mean ± standard error of the mean (SEM). Sample number (n) indicates the number of independent biological samples in each experiment. Sample numbers and experimental repeats are indicated in figures and figure legends. Statistical testing was carried out in the R software (Posit) or in GraphPad Prism using Student’s T-test, non-parametric Mann-Whitney test, or one-way or two-way analysis of variance (ANOVA) with Tukey’s post-hoc test as indicated in figure legends. Levels of significance are indicated as follows: p>0.05 (ns), p<0.05 (*), p<0.01 (**), p<0.001 (***), and p<0.0001 (****).

